# Claustrum–ACC reciprocal circuits modulate stress responses across acute and chronic states

**DOI:** 10.1101/2025.09.03.673892

**Authors:** Dan Liu, Zijun Liu, Gaojie Shao, Xiaoxuan Zhang, Haohao Hu, Shuai Chen, Xirong Xu, Jie Shao, Yiyun Qi, Zhiwei Ma, Qian Xiao, Yu Chen, Liping Wang, Fan Yang, Jie Tu

## Abstract

Anxiety is an evolutionarily conserved response that supports threat detection and survival, yet dysregulated stress responses can drive its progression into pathological states. Here, we identify the claustrum (CLA) as a central hub that links acute stress responses to chronic affective dysfunction through cell-type-specific circuit dynamics. Acute social defeat stress robustly activated the CLA and induced anxiety-like behaviors in mice. Multimodal profiling revealed transcriptional and functional plasticity within CLA neurons, with Egr2-expressing excitatory and Gad2-expressing inhibitory populations exhibiting distinct adaptations to acute versus chronic stress. We further delineate a reciprocal CLA–anterior cingulate cortex (ACC) circuit that bidirectionally regulates stress responses in which glutamatergic CLAEgr2 → ACC projections promoted anxiety-like behaviors, whereas top-down ACC → CLA^Gad2^inhibitory inputs constrained claustral output. Chronic stress induced hyperactivity in CLA^Gad2^ inhibitory neurons and reshaped excitatory neuron dynamics, leading to increased intrinsic excitability, local circuit imbalance, and ultimately, depression-like phenotypes. These findings define a dynamic claustrum-centered circuit that governs the transition from adaptive anxiety to maladaptive affective states through balanced Egr2–Gad2 activity, and show that chronic-stress driven disruption of this excitatory-inhibitory equilibrium impairs stress coping and drives pathological behavioral outcomes.

**Graphic summary:** Acute stress engages a claustrum (CLA)-centered network with activation of Egr2⁺ excitatory neurons, promoting anxiety-like behaviors. Chronic stress drives CLA^G142⁺^ neurons into a hyperactive state, enhancing local inhibition and suppressing excitatory output to induce depressive-like behaviors. A reciprocal CLA^E7182→^ACC feedforward and ACC→CLA^G142^ feedback circuit forms a state-dependent regulatory hub governing stress adaptation.

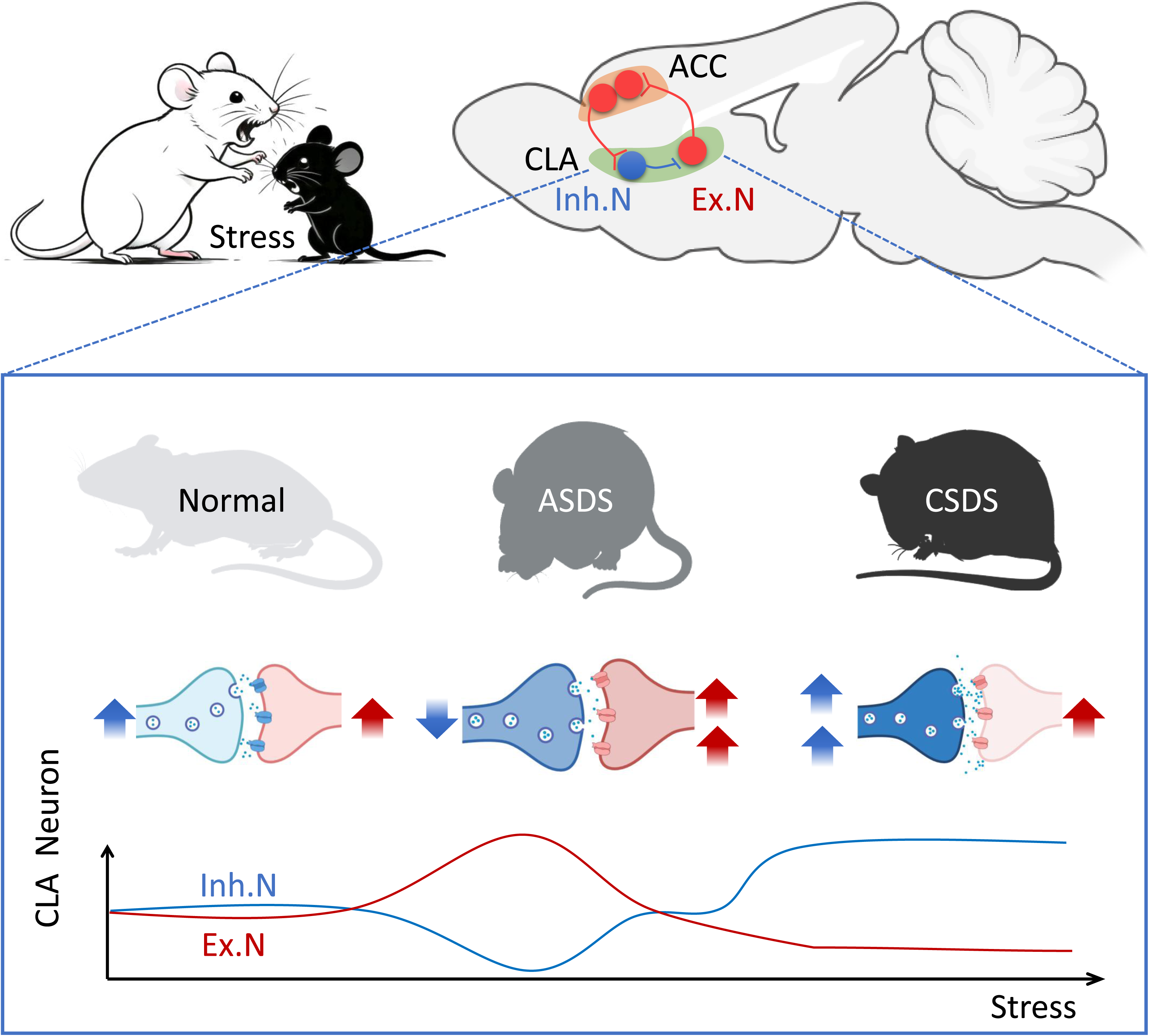

## INTRODUCTION

Anxiety is an evolutionarily conserved response that facilitates threat detection and promotes survival during acute stress.^1^ However, when stress is prolonged or excessive, this adaptive response can become maladaptive, driving the development of persistent anxiety and depression^2,3^. Chronic stress is known to disrupt neural activity, synaptic plasticity, and neurochemical signaling within emotion-related circuits^4–6^, yet a central unresolved question is how the brain dynamically regulates the transition from adaptive to pathological states.

Such stress regulation requires circuit mechanisms that actively balance excitatory and inhibitory activity to maintain behavioral stability under acute stress while preventing maladaptive escalation during chronic stress. Although this balance is fundamental to neural function, the specific circuits that implement this control, and how they dynamically reconfigure to govern transitions across stress states, remain largely unknown. A defined structure or region that can integrate acute stress signals and broadcast output to higher-order cortical regions to shape behavior has not yet been identified.

The claustrum (CLA), a densely interconnected subcortical structure, has emerged as a potential integrative node capable of coordinating cortical activity across diverse functional domains.^7,8^ Human imaging studies imaging studies indicate that the CLA is robustly engaged during acute stress, alongside regions such as the amygdala and thalamus.^9^ Through its extensive bidirectional connectivity with the cortex, the CLA is positioned to regulate arousal, attention, and behavioral responses to salient stimuli.^10,11^ In rodents, CLA activity has been linked to anxiety-like behaviours,^12^ and single-cell transcriptomic studies have identified distinct excitatory (vGluT1) and inhibitory (Gad2) neuronal populations within the CLA.^13–16^ However, it remains largely unknow how these cell types are organized into functional circuits that regulate stress responses, and whether such circuits dynamically reconfigure across stress conditions.

Here, we combine multimodal approaches, including a 3D Behavior Atlas, 9.4T functional magnetic resonance imaging (fMRI), fiber photometry, and single-nucleus RNA sequencing (snRNA-seq), to define the circuit and cellular mechanisms underlying stress adaptation in the CLA. We identify a claustrum-centered, state-dependent push–pull circuit that governs stress-related behavioral outcomes. Specifically, claustral Egr2⁺ excitatory neurons form a feedforward projection to the anterior cingulate cortex (ACC) that promotes anxiety-like behaviors, whereas inhibitory control is provided by intrinsic CLA^Gad2^ activity and is reinforced by a top-down ACC→CLA^Gad2^ feedback pathway. This opposing circuit architecture establishes a dynamic balance between excitatory drive and inhibitory restraint that enables adaptive responses to acute stress. Under chronic stress, this balance is progressively reconfigured, with a shift toward inhibitory dominance and altered excitability within claustral networks, ultimately leading to depression-like behaviors. Together, these findings reveal a claustrum-centered circuit mechanism that governs the transition from adaptive anxiety to maladaptive affective states.

## RESULTS

### Acute social defeat stress induces anxiety-like behavior without depressive phenotypes

To determine how acute stress alters behavioral states, we examined mice following acute social defeat stress (ASDS). To comprehensively characterize these alterations, we performed 3D behavioral tracking to quantify spontaneous behavior in an unbiased manner.^17^ Mice were individually placed in a cylindrical open field, and behavior was recorded for 15 minutes (Fig. 1A, Fig. S1A). A multi-view 3D animal motion-capture system^18^ enabled reconstruction of body posture and trajectories for fine-grained behavioral quantification (Fig. S1B).

**Figure 1.**
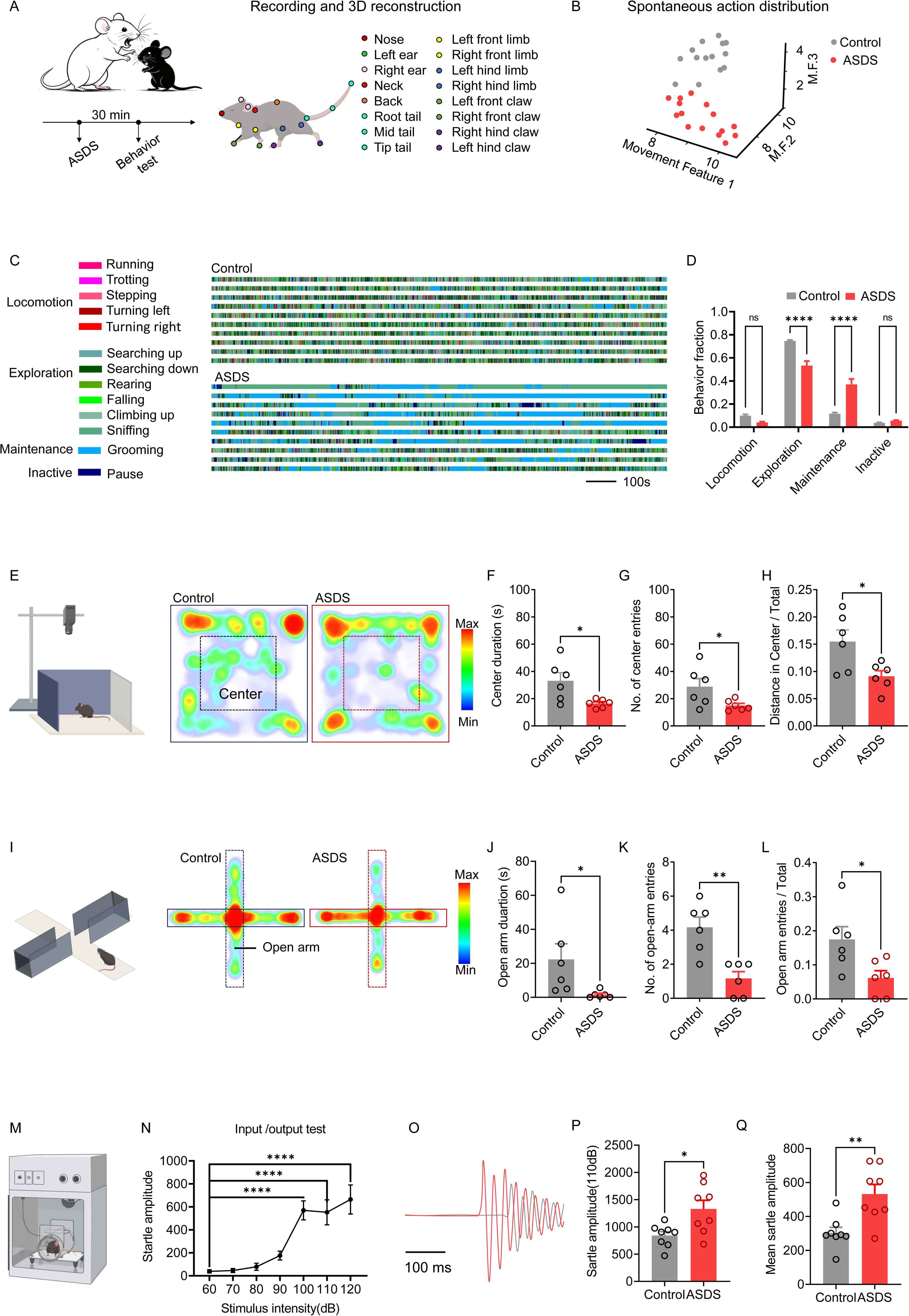
Anxiety-like behavior induced by acute social defeat stress (ASDS) (A) 3D behavioral tracking and pose reconstruction following ASDS. Left: Experimental timeline, in which mice received ASDS and rested for 30 min before behavioral testing. Right: Schematic showing 3D pose reconstruction, with colored markers indicating tracked anatomical landmarks (nose, ears, neck, back, tail segments, and limb/claw positions) for quantifying fine-grained social and locomotor behaviors. (B) Multivariate segregation of movement phenotypes in control and ASDS mice (n = 14–15/group). Three-dimensional embedding of behavior fractions by *t*-SNE shows distinct clustering of control (black) and ASDS (red) groups along Movement Features 1–3. Spatiotemporal feature space of 13 behavioral components was derived from video and 3D ethological skeleton plots to define body positions and behaviors. (C–D) Behavioral repertoire profiling reveals distinct ethological patterns in control and ASDS mice. (C) Ethogram of mouse behavior, categorized into four functional domains: Locomotion (Running, Trotting, Stepping, Turning left), Exploration (Searching down, Rearing, Falling, Climbing up), Maintenance (Grooming), and Inactive (Pause). (D) Proportion of time spent in each behavior quantified by automated 3D pose tracking (control, n = 15; ASDS, n = 14). Multivariate analysis revealed a significant main effect of behavior type (*F*₍₃,₁₀₈₎ = 299.6, *P* < 0.001) and a significant group × behavior interaction (*F*₍₃,₁₀₈₎ = 37.92, *P* < 0.001). Post-hoc Šidák tests showed that ASDS mice exhibited significantly reduced exploration (adjusted *P* < 0.0001) and increased maintenance (adjusted *P* < 0.0001) compared with controls. (E–H) Open-field test (OFT) in ASDS and control groups (n = 6/group). (E) Schematic showing the OFT paradigm, representative movement heatmaps of control and ASDS mice. (F–H) Quantification: (F) Time spent in the central zone (*t*(10) = 2.708, *P* = 0.0220). (G) Number of entries into the central zone (*t*(10) = 2.263, *P* = 0.0471). (H) Proportion of total activity in the central zone (*t*(10) = 2.720, *P* = 0.0216). Statistical analysis was performed using an unpaired two-tailed Student’s *t*-test. (I–L) Comparison of elevated plus maze (EPM) behavior between ASDS and control groups (n = 6/group). (I) Schematic showing the EPM paradigm, representative movement heatmaps of control and ASDS mice. (J–L) Quantification: (J) Open arm duration (*t*(10) = 2.251, *P* = 0.0481). (K) Number of open-arm entries (*t*(10) = 4.151, *P* = 0.0020). (L) Ratio of open-arm entries to total entries (*t*(10) = 2.620, *P* = 0.0256). Statistical analysis was performed using an unpaired two-tailed Student’s *t*-test. (M–N) Startle reflex in ASDS and control groups. (M) Schematic showing the startle reflex apparatus. (N) Validation of stable startle amplitude (n = 5/group, *F* = 14.52, *P*<0.0001). (O) Representative startle response waveforms. (P–Q) Quantification (n = 8/group): (P) Maximum startle amplitude (*t*(14) = 2.787, *P* = 0.0145). (Q) Mean startle amplitude (*t*(14) = 3.492, *P* = 0.0036). Statistical analysis was performed using an unpaired two-tailed Student’s *t*-test. Data are presented as mean ± SEM. \**P* < 0.05, \*\**P* < 0.01, \*\*\**P* < 0.001, \*\*\*\**P* < 0.0001, n.s. represents *P* > 0.05.

High-dimensional behavioral features embedded into a 3D coordinate space using t-SNE, where the spatial proximity of points reflects similarity in the original feature space (Fig. 1B). Clustering analysis, based on a previously established protocol,^17^ identified 13 discrete behavioral motifs, which were further grouped into four functional states: locomotion, exploration, maintenance, and inactivity (Fig. 1C, Fig. S1C–E). ASDS mice showed a marked reduction in exploratory behaviors as well as decreased locomotion, accompanied by increased maintenance and inactivity (Fig. 1D). Kinematic analysis revealed reduced body length, body height, and body rotation angles in ASDS mice relative to controls (Fig. S1F–H), indicating suppressed exploratory drive and increased behavioral constraint.

We next validated these findings using standard behavioral assays. In the open-field test (OFT), ASDS mice spent significantly less time in the center, made fewer center entries, and travelled shorter distances within the central zone compared to controls (Fig. 1E-H). Similarly, in the elevated plus maze (EPM), ASDS mice exhibited reduced open-arm exploration, reflected by decreased time, entries, and distance travelled in open arms (Fig. 1I-L), consistent with increased avoidance behavior.

To assess sensory reactivity, we measured acoustic startle responses and identified 110 dB as a reliable stimulus to evoke robust startle reflexes (Fig. 1M–N). Using this paradigm, ASDS mice displayed significantly increased peak and mean startle amplitudes relative to controls (Fig. 1O–Q), indicating heightened sensorimotor reactivity. ELISA measurements revealed elevated serum norepinephrine (NE) and corticosterone (CRF) levels in ASDS mice (Fig. S1I–K), confirming activation of physiological stress responses.

Importantly, ASDS did not induce depressive-like behaviors: sucrose preference and tail suspension tests showed no significant differences between groups (Fig. S1L–M). Together, these results define a distinct acute stress-induced behavioral state characterized by reduced exploratory drive, increased behavioral inhibition, heightened sensory responsiveness, and elevated physiological stress markers, without the emergence of depressive phenotypes.

### Distinct claustral neuronal subtypes are engaged by acute stress

To identify brain regions engaged during acute stress, we performed high-resolution resting-state imaging using 9.4T MRI^19^ in mice following ASDS. Consistent with previous human studies,^9^ whole-brain analysis revealed significant activation across multiple regions, with the CLA emerging as one of the most robustly activated structures (Fig. 2A-D). Functional connectivity analysis further indicated widespread reorganization of stress-related networks, including enhanced coupling between the CLA and cerebellum and nucleus accumbens shell (AcbSh), alongside reduced connectivity with contralateral CLA, primary somatosensory cortex (S1), and hippocampal CA3 (Fig. S2A-C). As a validation of signal specificity, chemogenetic activation of CLA^Egr2^ neurons elicited robust CLA activation in fMRI (Fig. S2D–E).

**Figure 2.**
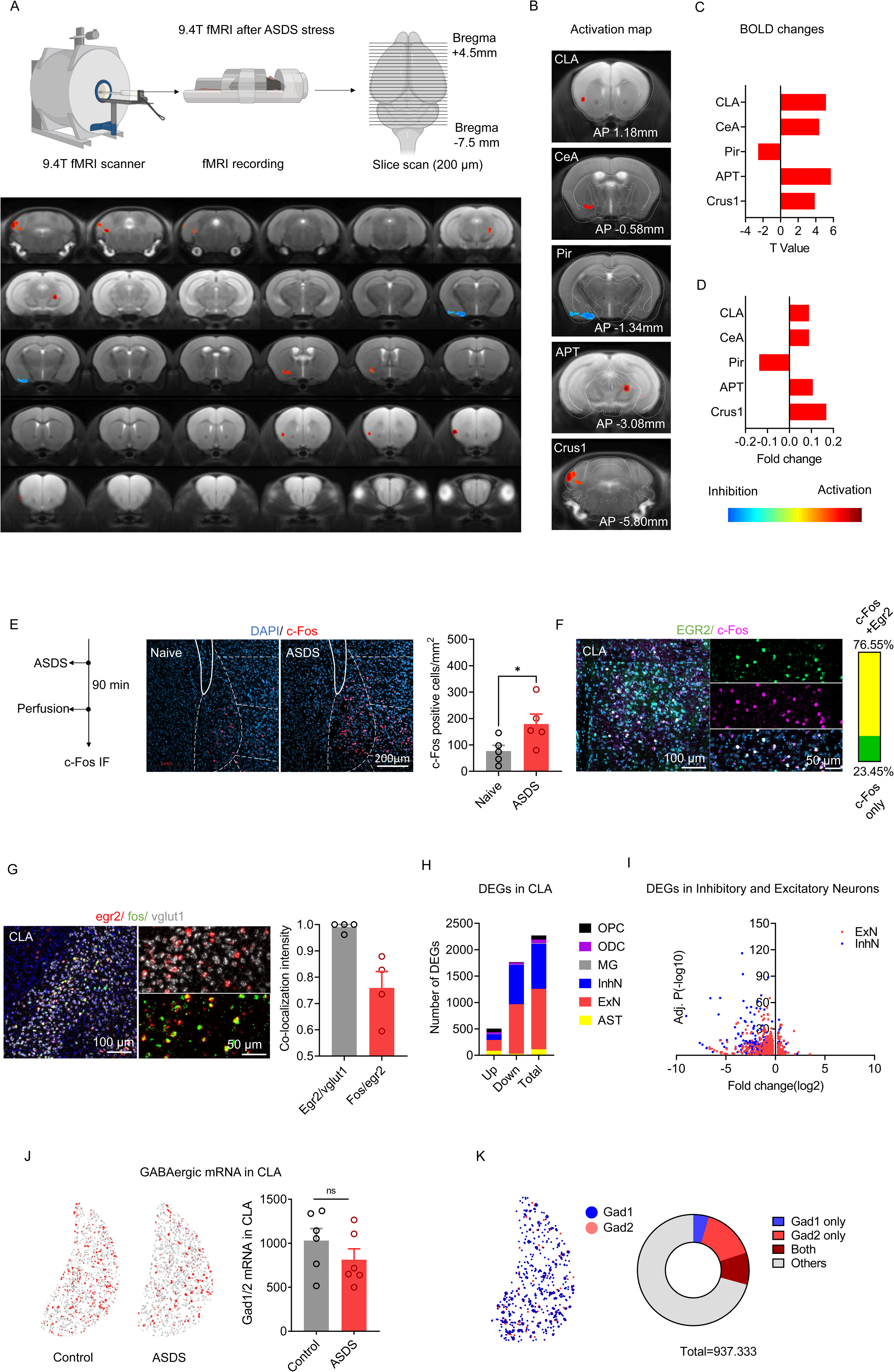
Changes in cellular and molecular profiles in the CLA of mice following ASDS. (A–D) (A) Schematic showing resting-state 9.4T fMRI acquisition under anesthesia following ASDS, with whole-brain fMRI Blood Oxygen Level Dependent (BOLD) signal analysis comparing ASDS and control groups (n = 10/group). (B) Merged coronal images from ASDS and control groups, illustrating whole-brain BOLD signal alterations and highlighting differentially activated brain regions corresponding to the brain atlas. (C) Statistical parametric mapping of BOLD signal T-values. (D) Functional connectivity (FC) ratios. Threshold: *P* < 0.01 with a 50-voxel extent. Statistical analysis was performed using an unpaired two-tailed Student’s *t*-test. (E) Immunofluorescence detection of c-Fos was performed in brain regions responsive to acute stress (n = 5/group), selected from the literature. Representative immunofluorescence images showing increased c-Fos activation in the CLA, with DAPI (cyan) labeling nuclei and c-Fos (red) labeling neuronal activation. Quantification of c-Fos-positive cell density, analyzed by an unpaired two-tailed Student’s t-test (*t*(8) = 2.337, *P* = 0.477). (F) Co-localization of c-Fos and EGR2 in the CLA of ASDS mice. Representative immunofluorescence staining and quantification showing that the majority of c-Fos^+^ activated neurons in the CLA co-express EGR2 in ASDS anxiety mouse model, analyzed by an unpaired two-tailed Student’s *t*-test (n = 9/group, *t*(16) = 32.36, *P <* 0.001). (G) Fos, egr2, vglut1 triple fluorescence *in situ* hybridization in CLAregion after ASDS. Representative confocal images showing fos (green), egr2 (red), vglut1 (grey) co-localization in CLA (n = 4/group). (H–I) Single-nucleus RNA sequencing (snRNA-seq) of the CLA following ASDS (ASDS → 60 min rest → CLA dissection → snRNA-seq, n = 2/group from 10 mice).(H) Quantification of differentially expressed genes (DEGs) across cell types in the CLA. (I) Cell-type enrichment analysis shows that the majority of DEGs originate from excitatory and inhibitory neurons. (J–K) Subtype analysis of inhibitory neurons in the CLA following ASDS. (J) Quantification of Gad1/2-positive cells revealed no significant change in density between control and ASDS groups (n = 6/group, *t*(10) = 0.5408, *P* = 0.6005). (K) Proportion of GABAergic neurons in the CLA brain region; Gad2-expressing neurons are the predominant subtype, accounting for 80.08%. Data are represented as mean ± SEM, \**P* < 0.05, ** *P <* 0.01, \*\*\**P* < 0.001, n.s.represents *P* > 0.05.

To independently validate these stress-induced network changes at cellular resolution, we performed whole-brain c-Fos mapping. ASDS mice exhibited significantly elevated c-Fos expression in multiple cortical, limbic, and neuroendocrine regions, including the ACC, mPFC, MD, CeA, BLA, PVN, and CA1 (Fig. S3A–L), whereas activity in regions such as the MO, NAc, VTA, and DRN remained unchanged (Fig. S3A-L). The CLA showed robust activation across both imaging and c-Fos analyses, supporting its recruitment as part of the stress-responsive network (Fig. S3).

We next examined the cellular basis of CLA activation. Combined *in situ* hybridization and immunofluorescence staining revealed that c-Fos activation after ASDS occurred predominantly in Egr2-positive (excitatory) neurons (Fig. 2F-G), indicating preferential recruitment of this neuronal subtype. To further characterize stress-induced cellular responses, we performed snRNA-seq to profile transcriptional changes across multiple cell types. Differential expression analysis revealed that stress-induced transcriptional changes were enriched in both excitatory and inhibitory neurons, as well as astrocytes (Fig. 2H). Despite their relative sparsity, inhibitory neurons exhibited a comparable number of differentially expressed genes relative to excitatory neurons (Fig. 2I), indicating disproportionately strong transcriptional plasticity within this population.

KEGG pathway analysis of these transcriptional changes revealed enrichment of metabolic and neurodegeneration-related pathways, including oxidative phosphorylation, Parkinson’s disease, and Huntington’s disease pathways, in both excitatory and inhibitory neurons (Fig. S2F–H), suggesting shared cellular stress-response program across neuronal subtypes.

To determine whether these changes reflected alterations in neuronal identity, we examined canonical marker expression within the CLA. Multiplex fluorescent *in situ* hybridization revealed no significant changes in the expression of GABAergic markers (Gad1 and Gad2) following ASDS (Fig. 2J), and Gad2 labeled the majority (∼95%) of CLA inhibitory neurons (Fig. 2K). Similarly, the number of cells expressing excitatory markers (including CaMKIIα, vGluT1, vGluT2 and Gnb4), inhibitory markers (Nos, Gad1 and Gad2), and other markers identified by sequencing or previous studies (Drd1, Drd2), remained comparable after ASDS, with expression levels stable across groups^20–21^ (Fig. S2I–J). These findings suggest that acute stress recruits the CLA at the network level and engages distinct neuronal subtypes, characterized by preferential activation of Egr2⁺ excitatory neurons and coordinated transcriptional responses across cell types, without altering neuronal identity.

### CLA^Gad2^ neurons exhibit state-dependent activity and bidirectionally control anxiety-like behavior

To determine how CLA^Gad2^ neurons respond to stress, we performed *in vivo* calcium recording in Gad2-Cre mice following control, acute (ASDS), or chronic stress (CSDS). GABAergic neurons in the CLA were selectively labeled by AAV-DIO-GCaMP6s, with immunostaining confirming high specificity (85.57% of GCaMP6s-expressing neurons co-localized with GABA, Fig. 3A–B).

**Figure 3.**
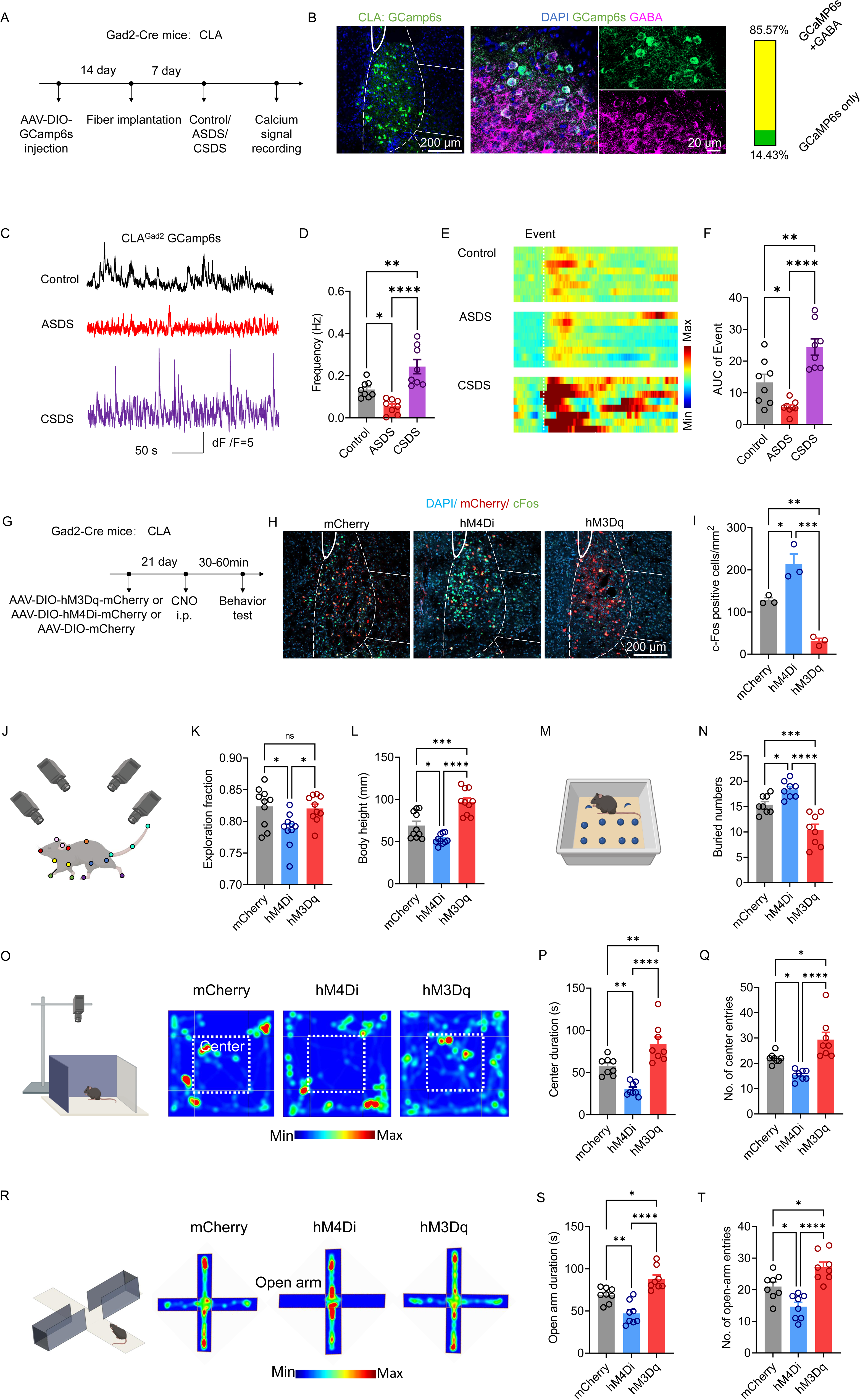
Regulatory role of CLA Gad2 neurons in anxiety-like behaviors. (A–F) Calcium signal recording of Gad2 neurons in the CLA. (A) Schematic diagram showing the virus injection and experimental workflow for calcium signal recording of Gad2 neurons in the CLA. (B) Schematic representation of GCaMP6s expression location and specificity in the CLA. (C) Representative traces of calcium signals in control, ASDS, and CSDS mice. (D) Quantification of firing frequency for events with amplitude > 2.91 and duration > 0.1 s, analyzed by Ordinary one-way ANOVA (n = 8/group, *F*(2, 21) = 18.5, *P* < 0.0001). (E) Heatmap of valid calcium transients from representative mice. (F) Quantification of the area under the curve (AUC) of valid calcium transients, analyzed by one-way ANOVA (n = 8/group, *F*(2, 21) = 19.92, *P* < 0.0001). (G–I) Chemogenetic modulation of CLA Gad2 neurons regulates anxiety-like behaviors. (G) Schematic showing the experimental workflow for chemogenetic manipulation of CLA Gad2 neurons. (H) Immunofluorescence staining for c-Fos was used to verify chemogenetic efficacy. (I) c-Fos density was analyzed by one-way ANOVA (n = 3/group, *F*(2,6) = 38.84, *P* = 0.0004). (J–L) 3D behavioral analysis revealed that inhibition of CLA Gad2 neurons decreased exploration time, analyzed by one-way ANOVA (n = 10/group, *F*(2, 27) = 5.655, *P*=0.0089). (L) Body height was increased by chemogenetic activation and decreased by inhibition, analyzed by one-way ANOVA (n = 10/group, *F*(2, 27) = 30.19, *P* < 0.0001). (M–N) (M) Schematic and results of the marble-burying test. (N) Quantification of the number of marbles buried by mice expressing mCherry (control), inhibitory hM4Di, or excitatory hM3Dq in the CLA, analyzed by one-way ANOVA (n = 8/group, *F*(2, 21) = 24.58, *P* < 0.0001). (O–Q) Representative heatmaps in the OFT (n = 8/group), quantification of time spent in the center zone analyzed by one-way ANOVA (*F*(2, 21) = 24.66, *P* < 0.0001), and number of entries into the center zone in control, ASDS, and CSDS mice(*F*(2, 21) = 16.16, *P* < 0.0001). (R–T) Representative heatmaps in the EPM (n = 8/group), quantification of time spent in open arms analyzed by one-way ANOVA (*F*(2, 21) = 22.48, *P* < 0.0001), and number of entries into open arms analyzed by one-way ANOVA (*F*(2, 21) = 18.09, *P* < 0.0001). Data are presented as mean ± SEM. \**P* < 0.05, \*\**P* < 0.01, \*\*\**P* < 0.001, \*\*\*\**P* < 0.0001, n.s. represents *P* > 0.05.

ASDS exposure led to a marked reduction in calcium transient frequency and area under the curve (AUC) in CLA^Gad2^ neurons compared to controls, indicating suppression of inhibitory activity under acute stress (Fig. 3C–F). In contrast, CSDS resulted in a significant increase in calcium transient frequency and AUC in CLA^Gad2^ neurons (Fig. 3C–F), indicating a reversal of activity dynamics across acute and chronic stress conditions. These results demonstrate that CLA^Gad2^ neurons exhibit bidirectional, state-dependent modulation across stress conditions.

To test whether CLA^Gad2^ neurons causally regulate anxiety-like behavior independent of stress exposure, we performed bidirectional chemogenetic manipulations. Gad2-Cre mice were injected with AAV-DIO-hM3Dq (activation), AAV-DIO-hM4Di (inhibition), or control virus in the CLA, and manipulation efficacy was confirmed by c-Fos immunostaining (Fig. 3G–I).

3D behavioral analysis revealed that inhibition of CLA^Gad2^ neurons significantly reduced exploration time and body height, indicating decreased exploratory drive (Fig. 3J–L). In contrast, activation of CLA^Gad2^ neurons increased body height relative to controls and partially rescued exploratory behavior compared to the inhibition group (Fig. 3J–L). Consistently, in the marble-burying assay, activation of CLA^Gad2^ neurons reduced marble-burying behavior, whereas inhibition increased it (Fig. 3M–N).

These effects were further validated using classical anxiety assays. In the OFT, activation of CLA^Gad2^ neurons increased time spent in the center and center entries, while inhibition produced the opposite effect (Fig. 3O–Q). Similarly, in the EPM, activation increased open-arm exploration, whereas inhibition reduced open-arm time and entries (Fig. 3R–T).

Together, these findings demonstrate that CLA^Gad2^ neurons are bidirectionally modulated by stress and exhibit state-dependent activity dynamics that causally regulate anxiety-like behavior. Acute stress suppresses their activity, whereas chronic stress induces a reversal of this pattern. Bidirectional manipulation of these neurons is sufficient to modulate behavior.

### Chronic stress drives hyperactivity of CLA^Gad2^ neurons to promote depressive-like behavior

We next examined whether chronic stress-induced alterations in CLA^Gad2^ neurons contribute to depressive-like behaviors. Consistent with the induction of depression-like states, CSDS mice exhibited a significant reduction in sucrose preference (Fig. 4A). To define the underlying cellular mechanisms, we performed electrophysiological recordings of CLA^Gad2^ neurons. CSDS, but not ASDS, robustly altered both synaptic input and intrinsic properties of CLA^Gad2^ neurons (Fig. 4B–M). Specifically, CSDS increased the amplitude and frequency of spontaneous excitatory postsynaptic currents (sEPSCs), indicating that Gad2-expressing neurons received enhanced glutamatergic synaptic drive, which may promote their excitability (Fig. 4C–E). In parallel, CLA^Gad2^ neurons displayed a marked increase in action potential output in response to current injection (Fig. 4F–G), reflecting heightened intrinsic excitability. This was accompanied by a depolarized action potential threshold and a significant reduction in rheobase current (Fig. 4I, M), whereas other electrophysiological properties, including action potential amplitude, half-width, and time constants, remained unchanged (Fig. 4H, J–L). Together, these results indicate that chronic stress induces a hyperexcitable state in CLA^Gad2^ neurons driven by enhanced excitatory input.

**Figure 4.**
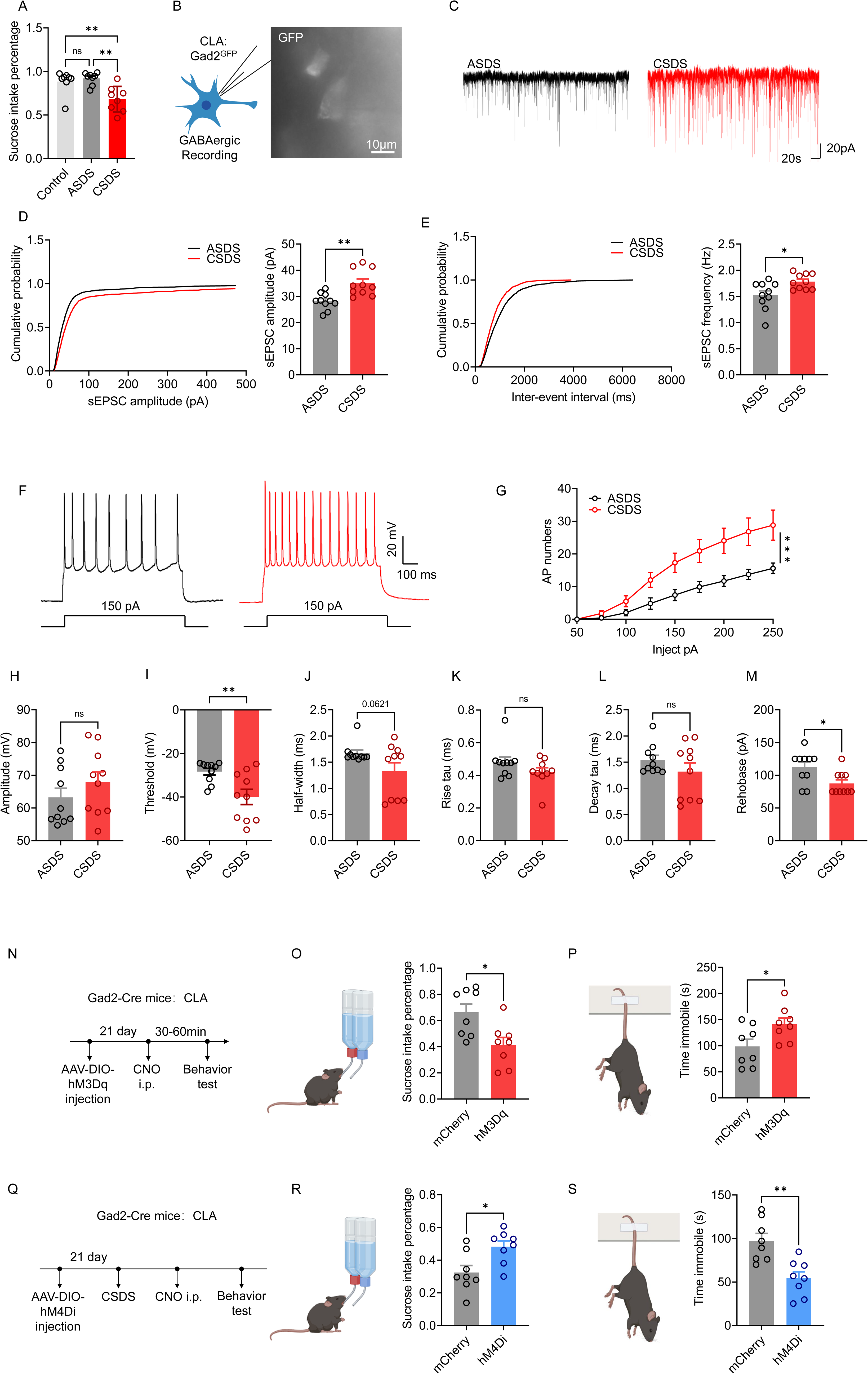
CLA Gad2 neurons respond to and regulate chronic social defeat stress-induced depressive-like behaviors. (A–E) CSDS enhances sEPSC activity in CLA Gad2 neurons and induces depressive - like behavior. (A) Compared with the ASDS group, the CSDS group successfully induced decreased sucrose preference in mice (n = 8/group, one-way ANOVA, *F* (2, 21) = 9.121, *P* = 0.0014). (B) Schematic showing the experimental workflow for electrophysiological recording of CLA GAD2 neurons following CSDS. (C) Representative traces of spontaneous excitatory postsynaptic currents (sEPSC). (D) Cumulative probability and quantification of sEPSC amplitude in CLA Gad2 neurons (n = 10/group), CSDS mice show significantly increased sEPSC amplitude, analyzed by an unpaired two-tailed Student’s t-test (*t*(18) = 3.618, *P* = 0.0017). (E) Cumulative probability and quantification of sEPSC frequency in CLA Gad2 neurons, CSDS mice show significantly increased sEPSC frequency, analyzed by an unpaired two-tailed Student’s t-test (*t*(18) = 2.671, *P* = 0.0156). (F–M) Altered intrinsic excitability and synaptic function in CLA Gad2 neurons following CSDS (n = 10/group). (F) Representative AP traces in CLA Gad2 neurons (150 pA injection) from ASDS and CSDS mice. (G) Input–output curve of AP numbers; CSDS mice show enhanced intrinsic excitability, analyzed by repeated-measures ANOVA (*F*(11, 198) = 7.030, *P* < 0.0001). (H) Quantification of action potential amplitude in ASDS and CSDS mice, analyzed by an unpaired two-tailed Student’s *t*-test (*t*(18) = 1.086, *P* = 0.2919). (I) Quantification of action potential threshold, showing a significantly more depolarized threshold in CSDS mice, analyzed by an unpaired two-tailed Student’s *t*-test (*t*(18) = 3.040, *P* = 0.0070). (J) Quantification of action potential half-width, analyzed by an unpaired two-tailed Student’s t-test (*t*(18) = 1.989, *P* = 0.0621). (K) Quantification of the rise time constant (rise tau) of action potentials, analyzed by an unpaired two-tailed Student’s t-test (*t*(18) =1.484, *P* = 0.1551). (L) Quantification of the decay time constant (decay tau) of action potentials, analyzed by an unpaired two-tailed Student’s t-test (*t*(18) = 1.176, *P* = 0.2550). (M) Quantification of the rheobase current, showing a significantly reduced rheobase in CSDS mice, analyzed by an unpaired two-tailed Student’s t-test (*t*(18) = 2.631, *P* = 0.0169). (N–P) Chemogenetic activation of CLA Gad2 neurons induces depressive-like behaviors(n = 8/group). (N) Experimental workflow for chemogenetic manipulation of CLA Gad2 neurons in Gad2-Cre mice. (O) Sucrose preference is reduced in hM3Dq mice, analyzed by an unpaired two-tailed Student’s t-test (*t*(18) = 2.924, *P* = 0.0111). (P) Immobility time in the tail suspension test is increased in hM3Dq mice, analyzed by an unpaired two-tailed Student’s t-test (*t*(18) = 2.374, *P* = 0.0324). (Q–S) Chemogenetic inhibition of CLA Gad2 neurons ameliorates CSDS-induced depressive-like behaviors(n = 10/group,). (Q) Experimental workflow for chemogenetic inhibition of CLA Gad2 neurons. (R) Sucrose preference is increased in hM4Di mice, analyzed by an unpaired two-tailed Student’s t-test (*t*(18) = 2.848, *P* = 0.0129). (S) Immobility time in the tail suspension test is decreased in hM4Di mice, analyzed by an unpaired two-tailed Student’s *t*-test (*t*(18) = 3.842, *P* = 0.0018). Data are presented as mean ± SEM. \**P* < 0.05, \*\**P* < 0.01, \*\*\**P* < 0.001, \*\*\*\**P* < 0.0001, n.s. represents *P* > 0.05.

To determine whether this hyperactivity causally contributes to depressive-like behavior, we performed bidirectional chemogenetic manipulations. Activation of CLA^Gad2^ neurons using hM3Dq recapitulated CSDS-like phenotypes, including reduced sucrose preference and increased immobility in the tail suspension test (TST) (Fig. 4N– P). Conversely, chemogenetic inhibition of CLA^Gad2^ neurons via hM4Di significantly ameliorated CSDS-induced depressive-like behaviors, restoring sucrose preference and reducing TST immobility (Fig. 4Q–S). Collectively, these findings demonstrate that chronic stress drives a hyperactive state of CLA^Gad2^ neurons through enhanced excitatory input, and that this state is both necessary and sufficient to promote depressive-like behavior, extending their state-dependent role from regulating anxiety under acute stress to driving maladaptive behavioral outcomes under chronic stress.

### CLA^Gad2^ neurons constrain excitatory activity within local CLA circuits

To assess the functional connectivity between inhibitory and excitatory populations in the CLA, we combined optogenetic activation of Gad2 neurons with whole-cell recordings in acute brain slices (Fig. S4A–D). Activation of Gad2 neurons reliably evoked inhibitory postsynaptic currents (IPSCs) in excitatory neurons, which were abolished by the GABAA receptor antagonist picrotoxin, confirming their GABAergic origin (Fig. S4E–G). In current-clamp mode, activation of Gad2 neurons increased the current required to evoke action potentials in excitatory neurons, indicating strengthened inhibitory control over excitatory output.

Consistent with the *in vitro* findings, fiber photometry recordings in awake mice confirmed that optogenetic activation of Gad2 neurons robustly suppressed calcium transients in vGluT1-positive excitatory neurons (Fig. S4H–K). During tail suspension stress, activation of Gad2 neurons attenuated stress-induced increases in excitatory neuronal activity (Fig. S4L–O), implying this inhibitory microcircuit dynamically constrains excitatory responses under acute stress conditions.

Together, these results demonstrate that acute stress induces a reorganization of inhibitory–excitatory interactions within the CLA, whereby strengthened synaptic inhibition onto excitatory neurons counterbalances network excitability. This local circuit mechanism provides a functional basis for their role in regulating stress-related activity.

### State-dependent reconfiguration of CLA^Egr^^2^ excitatory neurons governs anxiety and depressive behaviors

To determine how claustral excitatory neurons contribute to stress responses, we monitored the activity dynamics of CLA^Egr2^ neurons under acute and chronic stress conditions. *In vivo* calcium imaging in Egr2-Cre mice, with histological validation confirming ∼95% co-localization of GCaMP6s with CLA^Egr2^ neurons in the CLA (Fig. 5A–B), revealed that ASDS significantly increased both the frequency and AUC of calcium transients in these neurons relative to controls (Fig. 5C–F), confirming enhanced excitatory neuronal activity during acute stress.

**Figure 5.**
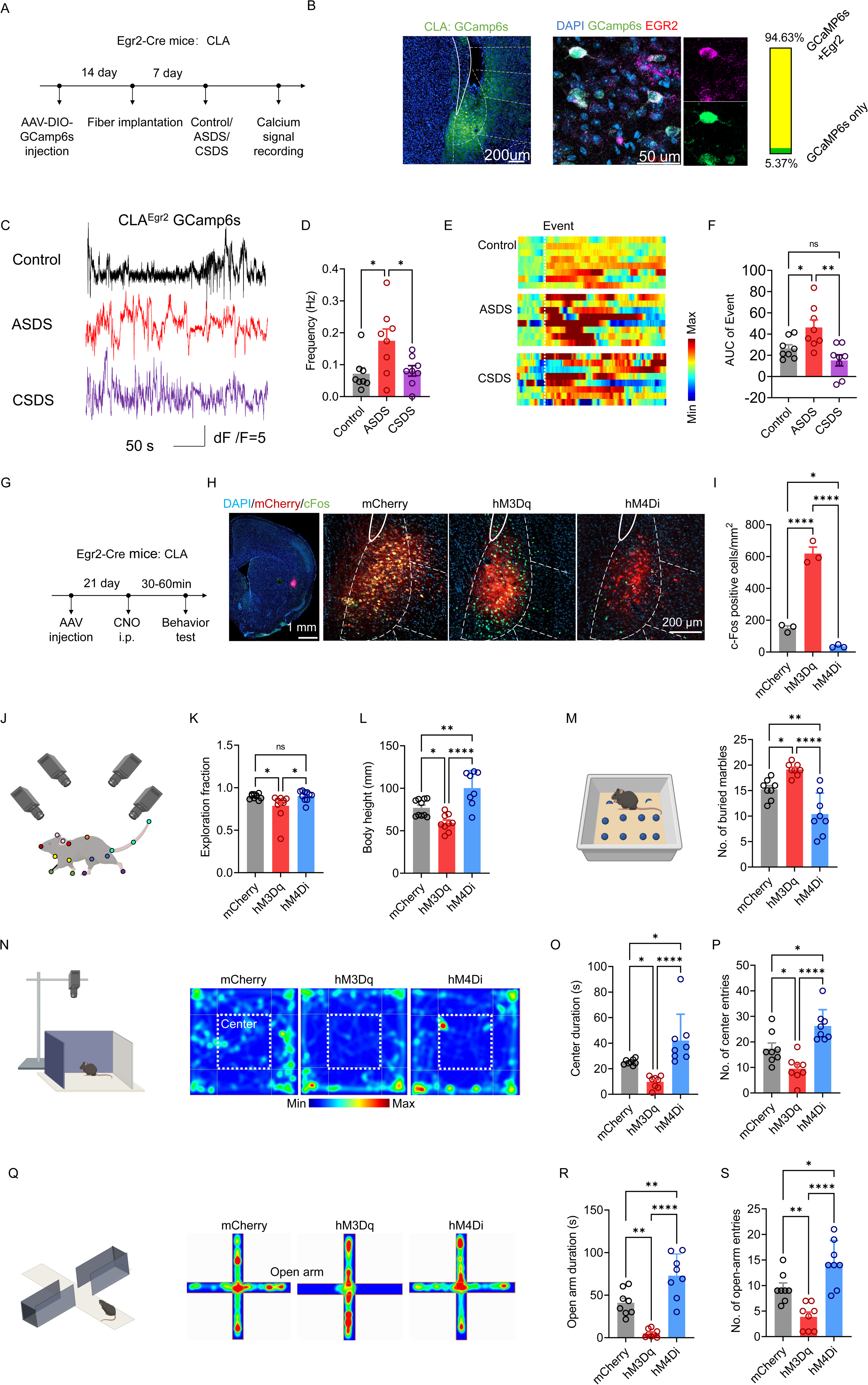
Social Defeat Stress Remodels CLA^Egr2^ Neuron Calcium Signaling and Chemogenetic Modulation of These Neurons Regulates Anxiety-Related Behaviors. (A–F) Chronic social defeat stress remodels calcium signaling in CLA^Egr2^ neurons. (A) Timeline of experiments: AAV-DIO-GCamp6s injection into CLA of Egr2-Cre mice, fiber implantation, Control/ASDS/CSDS stress, and *in vivo* calcium recording. (B) Immunofluorescence confirms GCamp6s (green) is expressed in CLA^Egr2^ neurons (red), with 94.63% co-localization; DAPI (blue) labels nuclei. (C) Representative calcium transient traces (*ΔF/F*) from Control, ASDS, and CSDS mice. (D) Quantification of calcium events frequency (n = 8/group, one-way ANOVA, *F* (2, 21) = 9.332, *P* = 0.0002). (E) Heatmap of calcium event amplitude dynamics, illustrating distinct activity patterns across Control, ASDS, and CSDS groups. (F) Quantification of area under the curve (AUC) of calcium events (n = 8/group, one-way ANOVA, *F* (2, 21) = 8.195, *P* = 0.0023). (G–I) Chemogenetic modulation of CLA^Egr2^ neurons regulates c-Fos expression. (G) Experimental timeline: AAV injection into CLA of Egr2-Cre mice, followed by CNO administration and behavioral testing after 21 days. (H) Representative immunofluorescence images showing mCherry (red), c-Fos (green), and DAPI (blue) in CLA. (I) Quantification of c-Fos-positive cells: hM3Dq activation significantly increases c-Fos expression, while hM4Di inhibition reduces it compared to mCherry controls (n = 3/group,one-way ANOVA, *F* (2, 6) = 150.9, *P* = 0.0001). (J–L) Chemogenetic modulation of CLA^Egr2^ neurons alters 3D behavioral analysis n = 8/group,. (J) Schematic showing the 3D behavioral analysis for spontaneous activity n = 8-10/group,. (K) Proportion of exploratory movements in 3D behavioral analysis (one-way ANOVA, *F* (2, 27) = 4.825, *P* = 0.0162). (L) Changes in body height in 3D behavioral analysis (one-way ANOVA, *F* (2, 24) = 18.43, *P* < 0.0001). (M) Schematic showing the marble-burying test after chemogenetic modulation of CLA^Egr2^ neurons. Reduced number of buried marbles in the marble burying test (n = 8/group, one-way ANOVA, *F* (2, 21) = 20.41, *P*<0.0001). (N–P) Chemogenetic modulation of CLA^Egr2^ neurons in the open field test (n = 8/group). (N) Schematic showing the open-field test and representative heatmaps of locomotor trajectories in mCherry, hM3Dq, and hM4Di groups. (O) Time spent in the central zone of the open field(one-way ANOVA, *F* (2, 21) = 13.79, *P* = 0.0001). (P) Number of entries into the central zone of the open field (one-way ANOVA, *F* (2, 21) = 16.45, *P* < 0.0001). (Q–S) Chemogenetic modulation of CLA^Egr2^ neurons in the elevated plus maze test(n = 8/group). (Q) Schematic showing the EPM test and representative heatmaps of locomotor trajectories in mCherry, hM3Dq, and hM4Di groups. (R) Time spent in the open arms of the EPM(one-way ANOVA, *F* (2, 21) = 30.66, *P* < 0.0001).(S) Number of entries into the open arms of the EPM (one-way ANOVA, *F* (2, 21) = 19.27, *P*<0.0001). Data are presented as mean ± SEM. \**P* < 0.05, \*\**P* < 0.01, \*\*\**P* < 0.001, \*\*\*\**P* < 0.0001, n.s. represents *P* > 0.05.

To further assess synaptic adaptations, we performed electrophysiological recordings of spontaneous excitatory postsynaptic currents (sEPSCs). While ASDS did not significantly alter glutamatergic input onto CLA^Egr2^ neurons, CSDS induced a marked reduction in both sEPSC amplitude and frequency (Fig. S5A–C), indicating weakened excitatory synaptic transmission under chronic stress.

We next examined the causal role of CLA^Egr2^ neurons in anxiety-like behavior using chemogenetic manipulation. Egr2-Cre mice were injected with AAV-DIO-hM3Dq (activation), AAV-DIO-hM4Di (inhibition), or control virus in the CLA, and manipulation efficacy was confirmed by c-Fos immunostaining (Fig. 5G–I). 3D behavioural analysis showed that activation of CLA^Egr2^ neurons significantly reduced exploration time and body height, whereas inhibition produced the opposite effect (Fig. 5J–L). Consistently, activation increased marble-burying behavior, while inhibition reduced it (Fig. 5M). In the OFT, activation decreased time spent in the center and centre entries, whereas inhibition exerted anxiolytic effects (Fig. 5N–P). Similarly, in the EPM, activation reduced open-arm exploration, whereas inhibition increased open-arm time and entries (Fig. 5Q–S). These findings demonstrate that activation of CLA^Egr2^ neurons is sufficient to drive anxiety-like behavior.

We then tested whether CLA^Egr2^ neurons contribute to chronic stress-induced depressive-like behaviors. Chemogenetic activation of CLA^Egr2^ neurons in CSDS mice significantly increased sucrose preference and reduced immobility in the TST, indicating an attenuation of depressive-like behaviors (Fig. S5D–F). Conversely, inhibition of CLA^Egr2^ neurons in naïve mice decreased sucrose preference and increased TST immobility (Fig. S5G–I), recapitulating core depressive-like phenotypes. These results indicate that CLA^Egr2^ neuronal activity is both necessary and sufficient to bidirectionally regulate depressive-like behavior.

Together, these findings demonstrate that CLA^Egr2^ neurons undergo state-dependent functional reconfiguration across acute and chronic stress conditions. Acute stress enhances excitatory neuronal activity to promote anxiety-like responses, whereas chronic stress is associated with reduced excitatory drive and a shift toward depressive - like states. In conjunction with the opposing dynamics observed in CLA^Gad2^ neurons, these results reveal a coordinated excitatory–inhibitory mechanism within the CLA that governs the transition from adaptive anxiety to maladaptive depression under prolonged stress.

### The CLA^Egr2^→ACC circuit mediates stress-related behavioral responses

Given the extensive efferent projections of the CLA to cortical regions, with the ACC as a major downstream target (Fig. S5J–N), and our observation of increased c-Fos activation in the ACC following ASDS, we next examined whether ACC excitatory neurons are engaged during acute stress. *In vivo* calcium imaging of CaMKIIα-positive activity specifically during social interaction (Fig. 6D–H). Representative traces and heatmaps showed pronounced activation of ACC excitatory neurons during stress exposure, and quantification of AUC confirmed a significant enhancement of activity in ASDS mice relative to baseline non-access periods (Fig. 6F–H), indicating dynamic recruitment of ACC neurons during acute stress.

**Figure 6.**
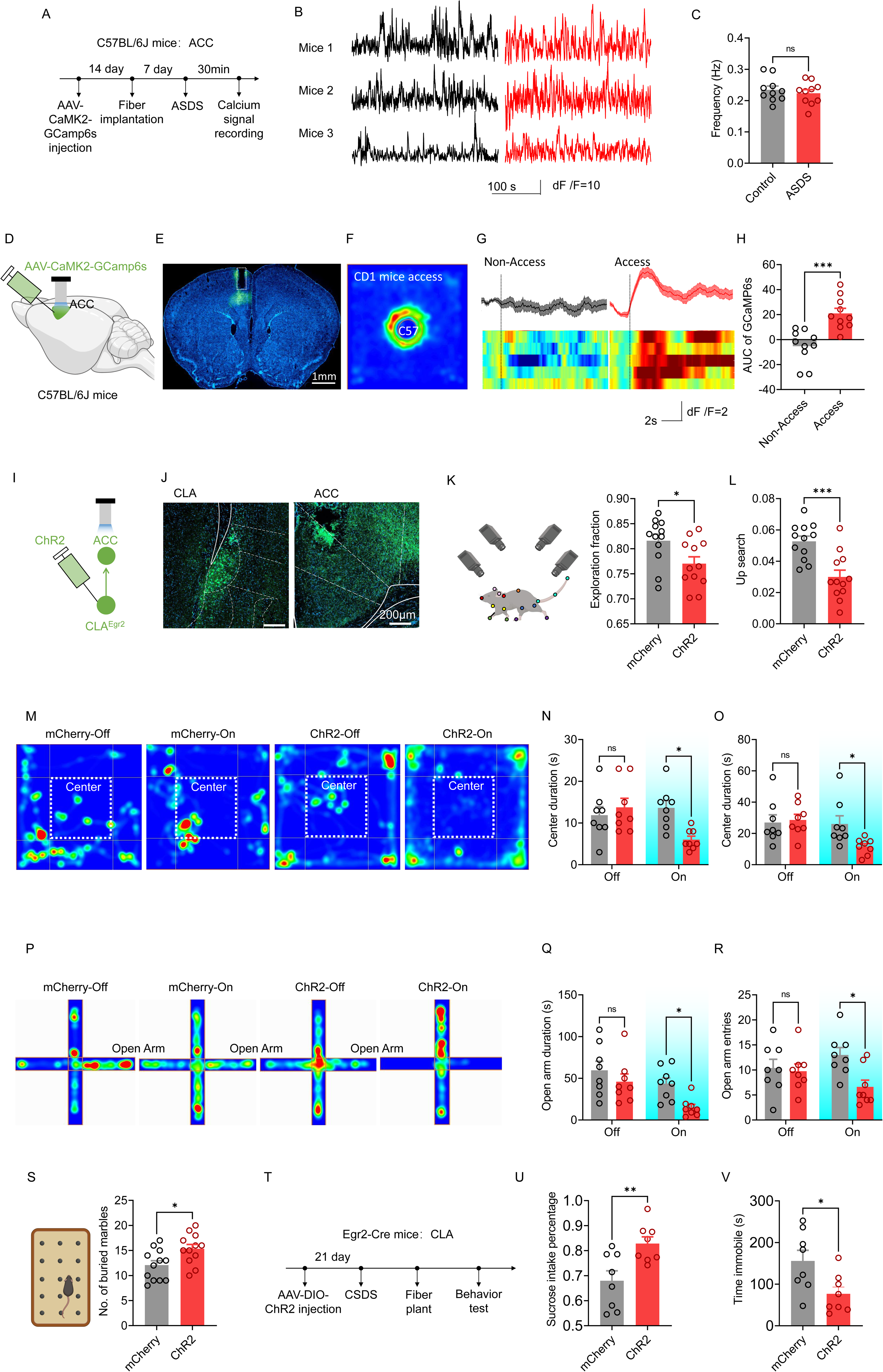
CLA^Egr2^→ACC Circuit Modulate Stress and Anxiety Behaviors. (A–H) Acute social defeat stress (ASDS) enhances calcium activity in ACC excitatory neurons during social interaction with CD1 mice. (A) Experimental timeline. (B) Representative calcium transient traces (*ΔF/F*) from ACC excitatory neurons in Control and ASDS mice. (C) Quantification of calcium event frequency in ASDS mice compared to Controls (n = 10/group, unpaired t-test, *t*(18) = 0.6528, *P* = 0.5221). (D) Schematic showing AAV-CaMKIIα-GCamp6s injection into the ACC of C57BL/6J mice for targeted labeling of excitatory neurons. (E) Representative coronal brain section confirming GCamp6s expression in the ACC. (F) Heatmap of ACC calcium activity during CD1 mice social defeat access. (G) Representative averaged *ΔF/F* traces and heatmaps showing enhanced calcium signals in ACC excitatory neurons during CD1 access. (H) Quantification of the area under the curve (AUC) of calcium responses demonstrates a significant elevation during social access (n = 10/group, unpaired *t*-test, *t*(18) = 4.467, P = 0.0003). (I–S) Optogenetic activation of the CLA^Egr2^→ACC circuit elicits anxiety-like behaviors. (I) Strategy for the CLA^Egr2^ → ACC projection-specific optogenetic activation. (J) Representative coronal brain section confirming ChR2 expression in the CLA. (K) Schematic showing the 3D behavioral analysis and quantification of the proportion of spontaneous exploratory movements following circuit activation (n = 12/group, unpaired *t*-test, *t*(22) =2.408, *P*=0.0249). (L) Up-search behavior is reduced upon circuit activation (n = 12/group, unpaired *t*-test, *t*(22) =4.169, *P*=0.0004). (M) Representative heatmaps of locomotor trajectories in the open field test. (N) Time spent in the central zone of the open field is reduced during optical activation (n = 8/group, Two-way ANOVA, *F* (1, 28) = 7.136, *P* = 0.0124). (O) The number of entries into the central zone of the open field is also reduced during light on (n = 8/group,Two-way ANOVA, *F* (1, 28) = 3.811, *P* = 0.0610). (P) Representative heatmaps of locomotor trajectories in the elevated plus maze (EPM) test. (Q) Time spent in the open arms of the EPM is reduced during optical activation (n = 8/group, Two-way ANOVA, *F* (1, 28) = 0.8393, *P* = 0.3674). R) The number of entries into the open arms of the EPM is also reduced during light on (n = 8/group, Two-way ANOVA, *F* (1, 28) = 3.622, *P* = 0.0674). (S) The number of buried marbles is increased following circuit activation (n = 12/group, unpaired t-test, *t*(22) =2.276, *P* = 0.0329). (T–V) Optogenetic activation of the CLA^Egr2^→ACC circuit alleviates depression-like behaviors following stress exposure. (T) Schematic showing the experimental timeline. (U) Sucrose preference test showing increased sucrose intake ratio upon optogenetic activation of this circuit(n = 8/group, unpaired t-test, *t*(14) = 3.095, *P* = 0.0079). (V) Tail suspension test showing reduced immobility time upon optogenetic activation of this circuit(n = 8/group, unpaired t-test, *t*(14) = 2.605, *P* = 0.0208). Data are presented as mean ± SEM. \**P* < 0.05, \*\**P* < 0.01, \*\*\**P* < 0.001, \*\*\*\**P* < 0.0001, n.s. represents *P* > 0.05.

To determine whether the CLA^Egr2^ → ACC projection mediates stress-related behavioral responses, we performed projection-specific optogenetic activation. Histological verification confirmed selective expression of ChR2 in CLA^Egr2^ somata and their terminals in the ACC (Fig. 6I, J). Optogenetic activation of the CLA^Egr2^→ ACC circuit recapitulated core anxiety-like phenotypes, as evidenced by reduced exploratory behavior in 3D behavioral analysis, including decreased spontaneous movement and up-search behavior (Fig. 6K, L), reduced center exploration in the open-field test (Fig. 6M–O), decreased open-arm exploration in the elevated plus maze (Fig. 6P–R), and increased marble-burying (Fig. 6S).

We next examined whether this circuit contributes to chronic stress-induced depressive-like behaviors. Consistent with the pro-resilience role of CLA^Egr2^ neurons, optogenetic activation of the CLA^Egr2^→ACC projection in CSDS-exposed mice significantly increased sucrose preference and reduced immobility time in the tail suspension test (Fig. 6T–V), indicating an attenuation of depressive-like phenotypes.

These findings demonstrate that the CLA^Egr^^2^→ACC circuit serves as a key output pathway through which claustral excitatory neurons influence stress-related behaviors, promoting anxiety-like responses under acute conditions while modulating behavioral outcomes under chronic stress.

### A reciprocal ACC→CLA^Gad2^ feedback circuit counterbalances CLA^Egr2^→ACC feedforward signaling to regulate stress responses

Given that the ACC is a major downstream target of the claustrum and exhibits reciprocal connectivity with it, we next investigated whether ACC projections provide feedback control onto CLA inhibitory neurons. Anterograde tracing experiments revealed that excitatory neurons in the ACC project to the CLA, with their synaptic boutons apposed around GABAergic interneurons (Figs. 7A–C). Retrograde labeling from CLA^Gad2^ neurons further identified ACC excitatory neurons as direct upstream inputs, establishing a monosynaptic ACC→CLA^Gad2^ feedback circuit (Fig. 7D–E).

**Figure 7.**
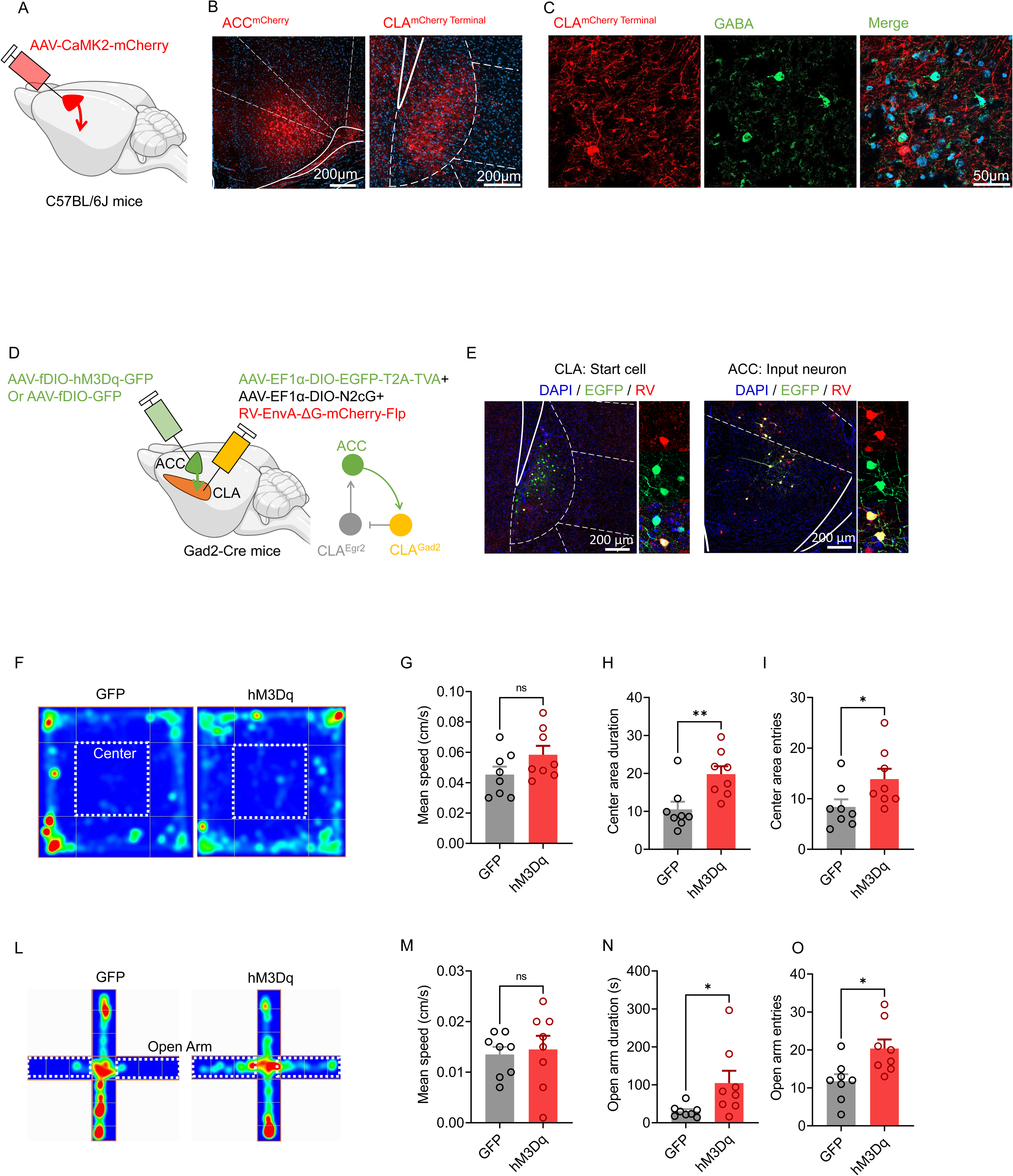
Activation of CLA^Egr2^→ACC and ACC→CLA^Gad2^ circuits alleviates stress-induced depressive- and anxiety-like behaviors. (A–C) Anterograde tracing reveals direct projections from the ACC to the CLA and axonal labeling surrounding CLA GABAergic neurons. (A) Schematic showing ACC circuit labeling. (B) Fluorescent protein expression in ACC cell bodies *in situ* and downstream terminals in the CLA. (C) Immunofluorescent staining showing colocalization of CLA terminals with GABA. (D–E) Labeled ACC→CLA^Gad2^ circuit. (D) Strategy for retrograde tracing from CLA Gad2-positive neurons to map the ACC → CLA^Gad2^ circuit. (E) Labeling of CLA-projecting neurons and their upstream input neurons. (F–I) Chemogenetic activation of ACC → CLA^Gad2^ circuit alleviates anxiety-like behaviors in the OFT (n = 8/group). (F) Representative heatmaps of animal locomotor trajectories. (G) Activation of ACC→CLA^Gad2^ does not alter locomotor speed in mice (unpaired t-test, *t*(14) = 1.665, *P* = 0.1181). (H) Increased time spent in the central zone of the OFT (unpaired *t*-test, *t*(14) = 3.170, *P* = 0.0068). (I) Increased number of entries into the central zone of the OFT (unpaired *t*-test, *t*(14) = 2.179, *P* = 0.0469). (L–O) Chemogenetic activation of ACC→CLA^Gad2^ alleviates anxiety-like behaviors in the elevated plus maze test (EPM). (L) Representative heatmaps of animal locomotor trajectories. (M) Activation of this circuit does not alter locomotor speed in mice (unpaired *t*-test, *t*(14) = 0.3257, *P* = 0.7495). (N) Increased time spent in the open arms of the elevated plus maze(unpaired t-test, *t*(14) = 2.260, *P* = 0.0403). (O) Increased number of entries into the open arms of the elevated plus maze (unpaired *t*-test, *t*(14) = 2.827, *P* = 0.0135). Data are presented as mean ± SEM. \**P* < 0.05, \*\**P* < 0.01, \*\*\**P* < 0.001, \*\*\*\**P* < 0.0001, n.s. represents *P* > 0.05.

To determine the functional role of this feedback pathway, we selectively activated ACC inputs to CLA^Gad2^ neurons using chemogenetic approaches. In the OFT, activation of the ACC→CLA^Gad2^ circuit did not affect locomotor speed but significantly increased time spent in the center and center entries, indicating reduced anxiety-like behavior (Fig. 7F–I). Similarly, in the EPM, circuit activation enhanced open-arm exploration without altering locomotion, as reflected by increased open-arm time and entries (Fig. 7L–O). These results demonstrate that ACC→CLA^Gad2^ feedback exerts a robust anxiolytic effect.

Together with the anxiogenic CLA^Egr2^→ACC feedforward pathway, these findings reveal a reciprocal circuit architecture in which feedforward excitatory signaling promotes anxiety-like responses, while feedback activation of inhibitory neurons provides counterbalancing control. This push–pull circuit motif establishes a dynamic excitatory–inhibitory regulatory system between the CLA and ACC, enabling flexible control of stress-induced behavioral states.

## DISCUSSION

Stress responses require coordinated neural activity to balance adaptive vigilance with emotional stability. Here, we show that the CLA contributes to this process through state-dependent changes in excitatory and inhibitory neuronal activity across acute and chronic stress conditions. Acute stress is associated with increased activity of CLA^Egr2^ excitatory neurons and suppression of CLA^Gad2^ inhibitory neurons, whereas chronic stress produces the opposite pattern, with reduced excitatory drive and enhanced inhibitory activity. At the circuit level, these opposing dynamics are reflected in reciprocal interactions between the CLA and ACC. Activation of the CLA^Egr2^→ACC projection drives anxiety, while reciprocal ACC→CLA^Gad2^ feedback suppresses it, supporting a push–pull organization linking claustral output with cortical feedback. Together, these findings support a model in which dynamic reconfiguration of excitatory–inhibitory balance within claustral–cortical circuits shapes behavioral responses to stress across states.

Under chronic stress, this balance is disrupted, with a shift toward inhibitory dominance and reduced circuit flexibility. Sustained hyperactivity of CLA^Gad2^ neurons, coupled with diminished Egr2⁺ activity, may bias this circuit toward a maladaptive state associated with depressive-like behavior. These observations suggest that state-dependent changes in excitatory–inhibitory balance within claustral circuits contribute to the emergence of distinct stress-related behavioral outcomes.

### Egr2–Gad2 microcircuit implements state-dependent push–pull control in the CLA

Research on the CLA has progressively advanced from population-level imaging to cellular and circuit-level analyses^12,22,23^, yet the intrinsic microcircuit organization underlying its role in stress regulation remains poorly defined. Here, we define a push–pull microcircuit formed by CLA^Egr2^ excitatory neurons and CLA^Gad2^ inhibitory neurons that dynamically controls behavioral responses across stress states.

Using multimodal behavioral, imaging, and molecular approaches, we found that both neuronal populations are engaged by stress but exhibit opposing activity dynamics depending on stress duration. CLA^Egr2^ neurons are preferentially activated under acute stress and suppressed under chronic stress, whereas CLA^Gad2^ neurons exhibit the opposite pattern, revealing a state-dependent reconfiguration of E/I balance within the CLA. Consistently, behavioral assays (3D and classical) revealed robust anxiety-like phenotypes following ASDS, accompanied by selective activation of the CLA in fMRI. Building on previous work showing that claustral excitatory neurons regulate anxiety without cell-type resolution,^12^ we combined multiplex *in situ* hybridization and snRNA sequencing to identify Egr2-expressing excitatory neurons and Gad2-expressing inhibitory neurons as the principal substrates of this response. *In vivo* calcium imaging further confirmed reciprocal activity trajectories across stress conditions, establishing an Egr2–Gad2 antagonistic motif within the claustral microcircuit.

Causal manipulations validate the functional significance of this organization. Activation of CLA^Egr2^ neurons increased anxiety-like behaviors, whereas inhibition reduced them. In contrast, activation of CLA^Gad2^ neurons exerted anxiolytic effects, whereas their inhibition promoted anxiety, identifying these neurons as a critical source of inhibitory constraint under physiological conditions. Notably, this protective function is inverted under chronic stress; CLA^Gad2^ neurons become pathologically hyperactive, leading to sustained suppression of excitatory output and the emergence of depressive-like behaviors.

Thus, the claustral Egr2–Gad2 microcircuit operates as a state-dependent gain-control system. Acute stress transiently biases the circuit toward excitatory dominance, promoting reversible anxiety-like responses, whereas chronic stress drives a shift toward inhibitory dominance, resulting in persistent depressive phenotypes. This transition from dynamic E/I balance to maladaptive inhibitory locking provides a local circuit mechanism linking stress duration to behavioral state change.

### The CLA–ACC bidirectional circuit constitutes a hierarchical control of emotional regulation

The claustral E/I microcircuit interfaces with long-range projections to form a hierarchical control system for emotional regulation. In this framework, the CLA–ACC axis operates as a bidirectional threat-processing loop in which feedforward excitation and feedback inhibition are dynamically coordinated. Specifically, the CLA^Egr2^→ACC projection amplifies anxiety-like responses, whereas the reciprocal ACC→CLA^Gad2^ pathway restores inhibitory balance, acting as a circuit-level constraint on stress escalation.

#### Anxiogenic circuit: CLA^Egr2^ → ACC

Excitatory neurons in the CLA densely innervate the ACC, and our data reveal that activation of the CLA^Egr2^→ACC pathway is sufficient to drive anxiety-like behaviors. This is consistent with previous studies showing that activation of claustral excitatory neurons promotes anxiety^24,25^, but extends these findings by identifying a functionally distinct Egr2^+^/vGluT1^+^ subpopulation that preferentially enhances cortical activity. Unlike the widely reported feedforward inhibitory influence of the claustrum on the prefrontal cortex via interneuron recruitment^12,25,26^, our results suggest that this projection can instead processing.

#### Anxiolytic circuit: ACC→CLA^Gad2^ feedback

In parallel, we identify a reciprocal ACC→CLA pathway that selectively engages inhibitory interneurons. Anatomical tracing confirms that ACC excitatory neurons project to Gad2-positive neurons within the CLA, consistent with prior evidence that cortical inputs recruit local GABAergic circuits.^27^ Functionally, activation of this pathway produces robust anxiolytic effects without altering locomotion, indicating that cortical feedback can directly suppress claustral output through inhibitory recruitment. This organization is further supported by prior studies showing that cortical inputs preferentially activate PV-positive interneurons in the claustrum.^16^ Given that Gad2-positive neurons encompass the major inhibitory populations within the CLA, including PV-positive interneurons,^13,28^ these findings support a model in which cortical inputs engage local inhibition to exert feedback control over claustral activity.

Our data further reveal that the CLA→ACC and ACC→CLApathways are mediated by distinct neuronal populations, forming a reciprocal but non-identical circuit architecture. Previous studies have described a disynaptic cross-hemispheric loop in which ACC neurons project to the contralateral CLA, recruit fast-spiking GABAergic interneurons, and in turn suppress cortical activity via an ACC → CLA → ACC inhibitory pathway.^29^ In line with these findings, our results and prior work^30^ indicate that ACC projections target both excitatory and inhibitory neurons within the CLA, while claustral outputs arise from distinct projection neuron populations. Although the precise multisynaptic organization of these pathways remains unresolved, in part because current viral strategies limit direct mapping of tri-synaptic connectivity, the available evidence supports a reciprocal, cell-type-specific feedback system linking the CLA and ACC.

Our data define two functionally opposing pathways—CLA^Egr^^2^→ACC as an anxiogenic feedforward projection, and ACC→CLA^Gad2^ as an anxiolytic feedback circuit—that form a tripartite reciprocal motif (Egr2–ACC–Gad2). This architecture provides a circuit-level implementation of stress allostasis, in which feedforward excitation and feedback inhibition are dynamically balanced under acute conditions but become destabilized with prolonged stress exposure

### The claustrum gates stress allostasis and the transition from adaptive anxiety to depression

A central finding of this study is that CLA neurons exhibit distinct and state - dependent regulatory patterns across anxiety- and depression-like behaviors, highlighting a critical role for the CLA in the transition from adaptive stress responses to affective pathology. ASDS shifts the circuit toward excitatory dominance through activation of CLA^Egr2^ neurons and suppression of CLA^Gad2^ neurons, promoting reversible anxiety-like behaviors. In contrast, CSDS induces persistent hyperactivity of CLA^Gad2^ neurons, reduces the E/I ratio, and drives depressive-like phenotypes. Bidirectional manipulations further demonstrate causality: activation of Gad2 neurons or inhibition of Egr2 neurons induces depression-like behaviors, whereas the opposite manipulations restore behavioral flexibility.

This view expands the classical stress-circuit framework centered on the amygdala. Although the amygdala is widely regarded as a core hub for acute stress responses, the CLA has traditionally been linked to sensory integration, conscious processing, and sleep–wake regulation^31–33^. However, its extensive connectivity with the amygdala, together with evidence of coactivation with the basolateral amygdala (BLA) during acute social defeat stress,^34^ supports the idea that the CLA participates in a broader emotional salience network. Our optogenetic and chemogenetic data further indicate that behavioral output is governed by the relative dominance of CLA^Egr2^ and CLA^Gad2^ neuronal activity, positioning the CLA as an integrative hub linking sensory, cognitive, and affective processes.

Beyond local microcircuit dynamics, MRI analyses reveal stress-induced remodeling of CLA-centered networks, disrupting the coordination of sensory, reward, and emotional systems. Increased functional connectivity between the CLA and the nucleus accumbens shell (AcbSh) and cerebellum may contribute to maladaptive reward processing^35^ and motor compensation,^36^ whereas reduced coupling with the hippocampus and cortex likely impairs memory integration and sensory processing.^37^ In addition, the emergence of negative correlations between ipsilateral and contralateral CLA activity suggests a disruption of interhemispheric inhibitory balance. These circuit-level alterations become progressively irreversible with repeated stress exposure, consistent with a transition from adaptive plasticity to pathological circuit reorganization.

Under chronic stress conditions, these changes are reinforced, with dominant activation of CLA^Gad2^ neurons driving depressive-like behaviors. Sustained hyperactivity of Gad2 neurons alongside reduced activity of Egr2 neurons is consistent with previous findings that chronic stress weakens excitatory CLA output to the prelimbic cortex via dynorphin/κ-opioid receptor signaling.^14^ This persistent inhibitory bias likely suppresses claustral excitatory drive more broadly, contributing to anhedonia and depressive-like phenotypes. In this context, prolonged hyper-defensive responses may not simply persist, but instead collapse into a hypo-flexible and pathologically constrained network state characterized by reduced E/I balance and impaired emotional adaptation.

Collectively, the CLA functions as a dynamic regulator of stress adaptation, in which both local microcircuit balance and long-range ACC–CLA interactions jointly shape behavioral outcomes. Acute stress transiently enhances excitability and engages top-down regulatory feedback, whereas chronic stress drives sustained inhibitory dominance and disrupts this regulatory balance, leading to maladaptive plasticity. In this context, CLA^Gad2⁺^ neurons function as a tonic inhibitory ‘brake’ that must be tightly regulated; once this constraint becomes excessive, emotional homeostasis breaks down, promoting depressive-like behaviors. Together, these findings support a unifying model in which the CLA operates as a state-dependent gain-control system embedded within a distributed ACC–CLA circuit, integrating local E/I balance, long-range interactions, and network-level dynamics to gate stress allostasis and its transition to pathology.

### Study limitations and translational potential

Despite these advances, several key limitations and open questions remain. Viral targeting of Gad2-positive neurons may extend beyond the claustrum, constraining cell-type and regional specificity, while the mechanisms underlying interhemispheric claustral coordination remain unresolved. At the cellular level, intrinsic membrane properties, synaptic inhibition, or neuromodulatory signaling require further validation. At the circuit level, although the CLA–ACC axis represents a core motif, the broader network architecture linking the CLA with the amygdala, striatum, thalamus, and other stress-related regions remains incompletely defined, as does the relative contribution of local interneurons versus long-range GABAergic projections. In addition, the progressive loss of reversibility observed under chronic stress suggests a critical transition window between adaptive compensation and pathological fixation, the timing and mechanisms of which remain to be further determined.

Addressing these challenges will require next-generation intersectional genetic tools, *in vivo* electrophysiological recordings during stress, and integrative multi-omics analyses across defined cell types and circuit nodes. Importantly, these findings also highlight the translational potential of state-dependent claustral circuit dynamics. Modulation of the CLA–ACC axis, particularly ACC→CLA feedback, may provide a strategy to restore disrupted inhibitory control in stress-related disorders. More broadly, targeting claustral E/I balance may offer a circuit-informed approach for treating comorbid anxiety and depression.

## STAR METHODS

### KEY RESOURCES TABLE

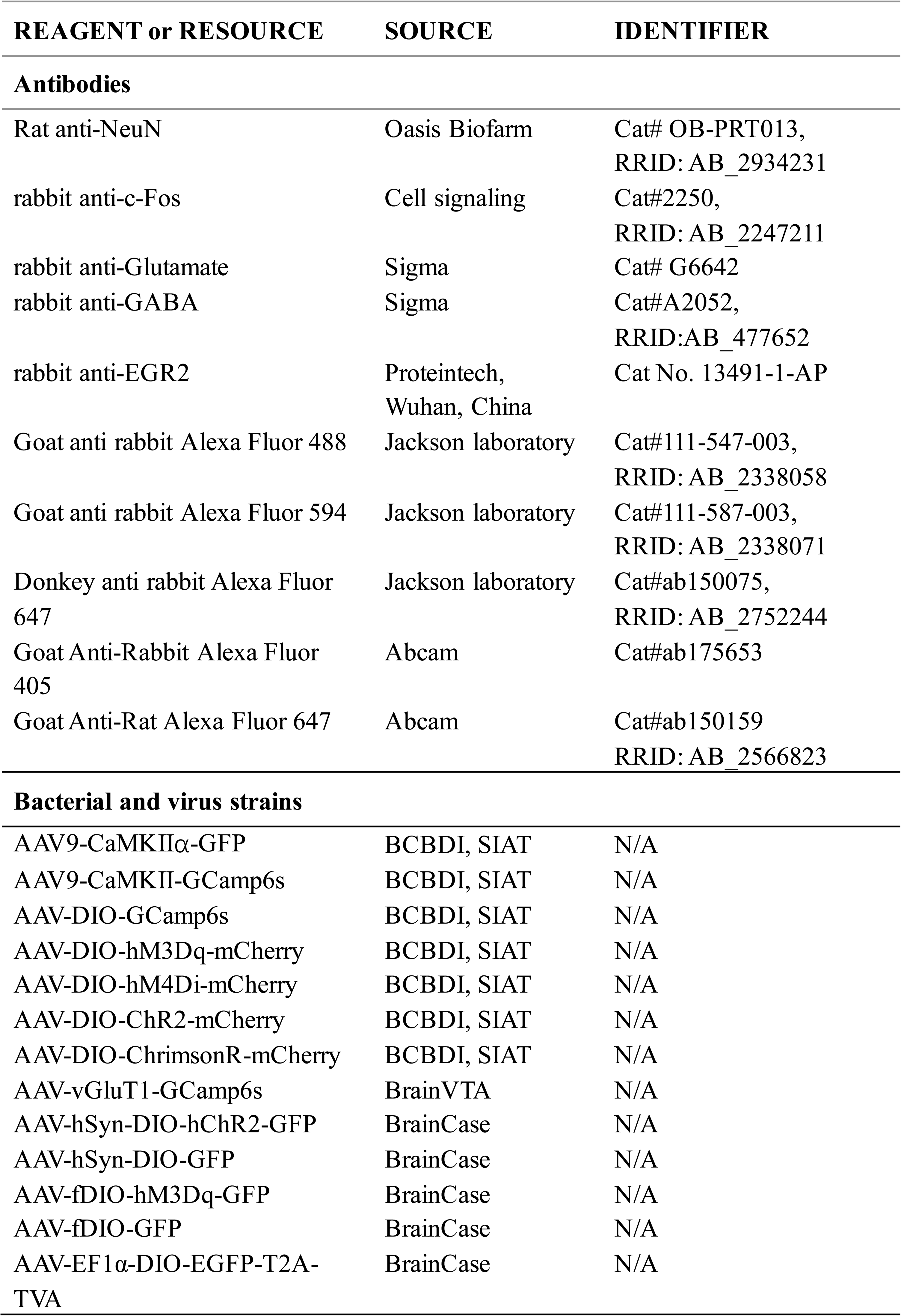
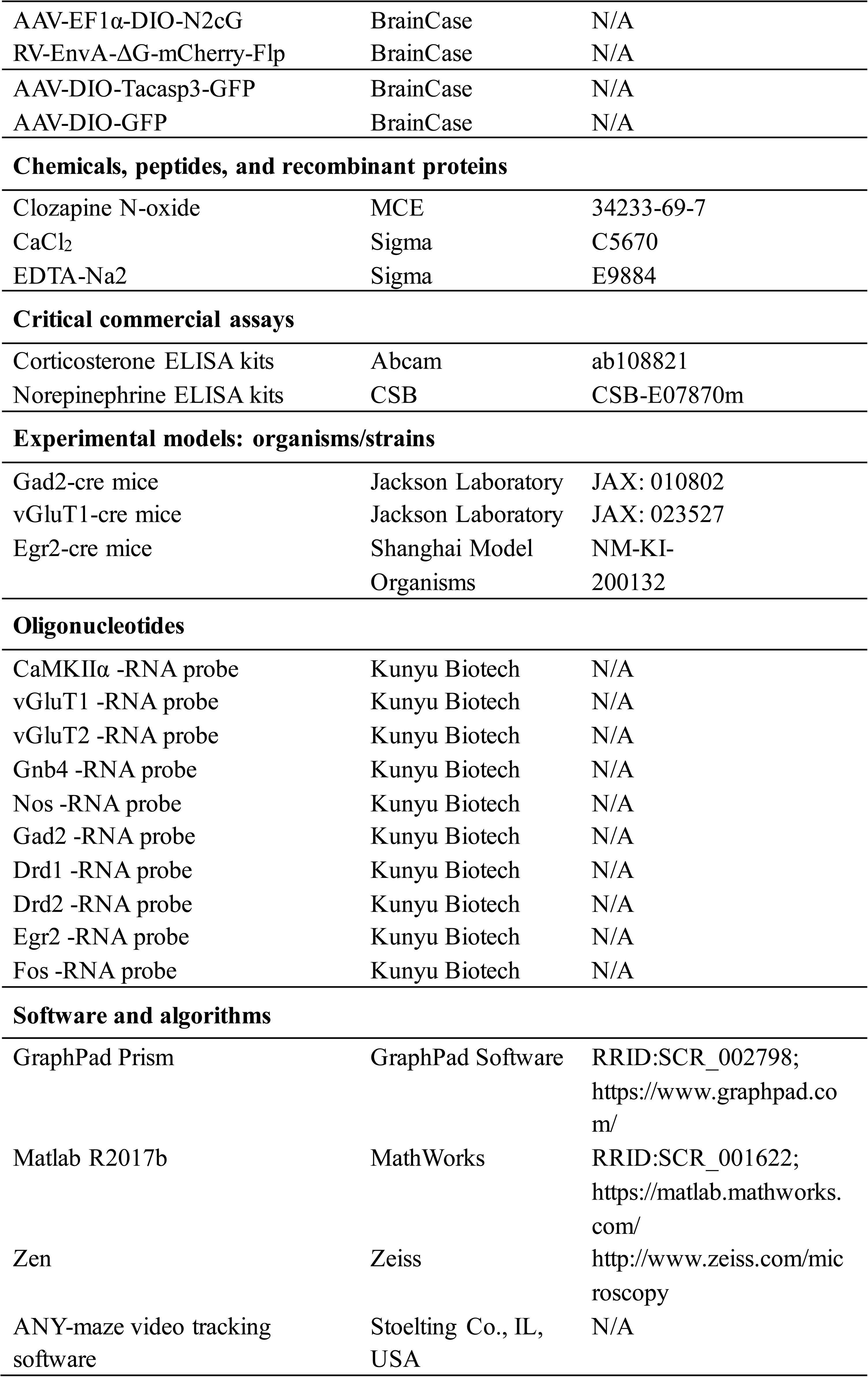
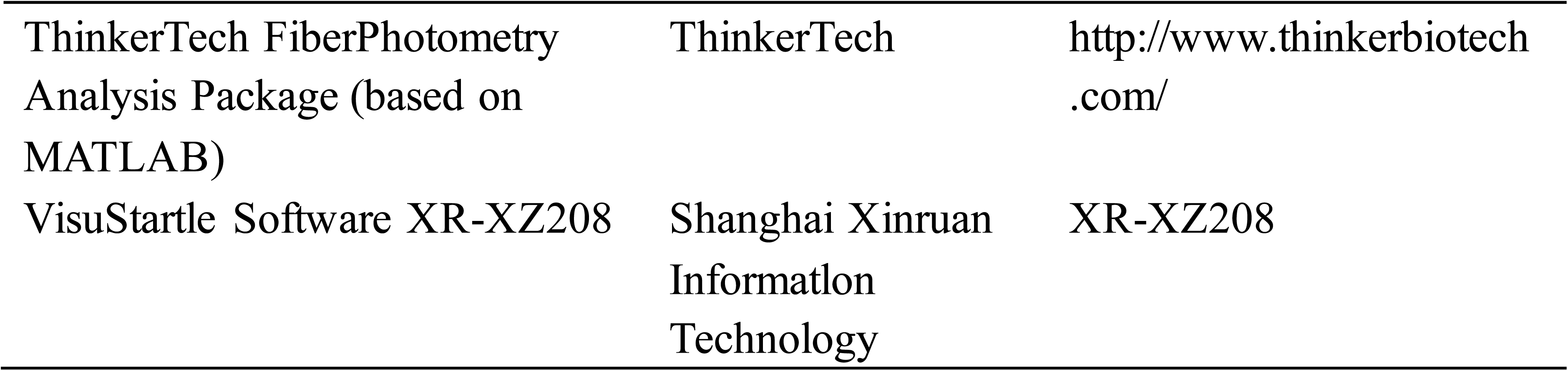

2 KEY RESOURCES TABLE
| REAGENT or RESOURCE | SOURCE | IDENTIFIER |
| --- | --- | --- |
| **Antibodies** |  |  |
| Rat anti-NeuN | Oasis Biofarm | Cat# OB-PRT013,<br>RRID: AB_2934231 |
| rabbit anti-c-Fos | Cell signaling | Cat#2250,<br>RRID: AB_2247211 |
| rabbit anti-Glutamate | Sigma | Cat# G6642 |
| rabbit anti-GABA | Sigma | Cat#A2052,<br>RRID:AB_477652 |
| rabbit anti-EGR2 | Proteintech,<br>Wuhan, China | Cat No. 13491-1-AP |
| Goat anti rabbit Alexa Fluor 488 | Jackson laboratory | Cat#111-547-003,<br>RRID: AB_2338058 |
| Goat anti rabbit Alexa Fluor 594 | Jackson laboratory | Cat#111-587-003,<br>RRID: AB_2338071 |
| Donkey anti rabbit Alexa Fluor 647 | Jackson laboratory | Cat#ab150075,<br>RRID: AB_2752244 |
| Goat Anti-Rabbit Alexa Fluor 405 | Abcam | Cat#ab175653 |
| Goat Anti-Rat Alexa Fluor 647 | Abcam | Cat#ab150159<br>RRID: AB_2566823 |
| **Bacterial and virus strains** |  |  |
| AAV9-CaMKIIα-GFP | BCBDI, SIAT | N/A |
| AAV9-CaMKII-GCamp6s | BCBDI, SIAT | N/A |
| AAV-DIO-GCamp6s | BCBDI, SIAT | N/A |
| AAV-DIO-hM3Dq-mCherry | BCBDI, SIAT | N/A |
| AAV-DIO-hM4Di-mCherry | BCBDI, SIAT | N/A |
| AAV-DIO-ChR2-mCherry | BCBDI, SIAT | N/A |
| AAV-DIO-ChrimsonR-mCherry | BCBDI, SIAT | N/A |
| AAV-vGluT1-GCamp6s | BrainVTA | N/A |
| AAV-hSyn-DIO-hChR2-GFP | BrainCase | N/A |
| AAV-hSyn-DIO-GFP | BrainCase | N/A |
| AAV-fDIO-hM3Dq-GFP | BrainCase | N/A |
| AAV-fDIO-GFP | BrainCase | N/A |
| AAV-EF1α-DIO-EGFP-T2A-TVA | BrainCase | N/A |
| AAV-EF1α-DIO-N2cG | BrainCase | N/A |
| RV-EnvA-ΔG-mCherry-Flp | BrainCase | N/A |
| AAV-DIO-Tacasp3-GFP | BrainCase | N/A |
| AAV-DIO-GFP | BrainCase | N/A |
| **Chemicals, peptides, and recombinant proteins** |  |  |
| Clozapine N-oxide | MCE | 34233-69-7 |
| CaCl<sub>2</sub> | Sigma | C5670 |
| EDTA-Na<sub>2</sub> | Sigma | E9884 |
| **Critical commercial assays** |  |  |
| Corticosterone ELISA kits | Abcam | ab108821 |
| Norepinephrine ELISA kits | CSB | CSB-E07870m |
| **Experimental models: organisms/strains** |  |  |
| Gad2-cre mice | Jackson Laboratory | JAX: 010802 |
| vGluT1-cre mice | Jackson Laboratory | JAX: 023527 |
| Egr2-cre mice | Shanghai Model<br>Organisms | NM-KI-200132 |
| **Oligonucleotides** |  |  |
| CaMKIIα -RNA probe | Kunyu Biotech | N/A |
| vGluT1 -RNA probe | Kunyu Biotech | N/A |
| vGluT2 -RNA probe | Kunyu Biotech | N/A |
| Gnb4 -RNA probe | Kunyu Biotech | N/A |
| Nos -RNA probe | Kunyu Biotech | N/A |
| Gad2 -RNA probe | Kunyu Biotech | N/A |
| Drd1 -RNA probe | Kunyu Biotech | N/A |
| Drd2 -RNA probe | Kunyu Biotech | N/A |
| Egr2 -RNA probe | Kunyu Biotech | N/A |
| Fos -RNA probe | Kunyu Biotech | N/A |
| **Software and algorithms** |  |  |
| GraphPad Prism | GraphPad Software | RRID:SCR_002798;<br>https://www.graphpad.com/ |
| Matlab R2017b | MathWorks | RRID:SCR_001622;<br>https://matlab.mathworks.com/ |
| Zen | Zeiss | http://www.zeiss.com/microscopy |
| ANY-maze video tracking software | Stoelting Co., IL,<br>USA | N/A |
| ThinkerTech FiberPhotometry<br>Analysis Package (based on<br>MATLAB) | ThinkerTech | http://www.thinkerbiotech.com/ |
| VisuStartle Software XR-XZ208 | Shanghai Xinruan<br>Information<br>Technology | XR-XZ208 |

#### Resource availability

##### Lead contact

Further information and requests for resources and reagents should be directed to and will be fulfilled upon reasonable request by lead contact, Jie Tu.

#### Materials availability

The transgenic mouse lines Egr2-Cre, vGluT1-Cre and Gad2-Cre mice are available upon reasonable request after signing a material transfer agreement with the Shenzhen Institutes of Advanced Technology, Chinese Academy of Sciences.

#### Experimental model and subject details

##### Animals

All experimental procedures were approved by the Institutional Animal Care and Use Committee (IACUC Protocol #SIAT-IACUC-210201-NS-LD-A1539) of Shenzhen Institutes of Advanced Technology of the Chinese Academy of Sciences. Male C57BL/6J and CD1/ICR mice were purchased from the Zhejiang Weitong Lihua Company. Transgenic mice expressing Cre recombinase under the control of the *Egr2* (Egr2-Cre, Stock # NM-KI-200132), *Slc17a7* (vGluT1-Cre, Stock #023527) or *Gad2* (Gad2-Cre, Stock #010802) promoters were obtained from The Jackson Laboratory. All transgenic lines were maintained on a C57BL/6J background. Mice were group-housed (3−5 per cage) in a temperature- and humidity-controlled environment under a 12-hour light/dark cycle (lights on at 07:00), with ad libitum access to food and water. Experimental cohorts consisted of age-matched littermates (8−12 weeks old) randomly assigned to experimental or control groups. To minimize circadian variability, all behavioral testing was conducted during the light phase. Efforts were made to minimize animal suffering, including environmental enrichment and post-surgical analgesia where applicable.

##### Behavioral tests

###### Acute social defeat stress (ASDS)

ASDS was conducted using a modified version of the protocol established by Golden et al.^38^ Male CD1 mice (8−12 weeks old) were pre-screened for consistent aggressive behavior, defined as an attack latency of less than 30 seconds and a minimum of 10 attacks per session for three consecutive days. Only animals meeting these criteria were used as aggressors. Experimental male C57BL/6J mice (8−10 weeks old) were individually introduced into the home cage of a screened aggressor and subjected to 10 physical attacks over a 10−15-minute period. Upon submissive posturing (defined as immobility or freezing within 3 seconds after each attack), the experimental mouse was immediately separated from the aggressor using a perforated transparent divider, allowing continued sensory contact (visual, olfactory, and auditory) while preventing further physical interaction and injury.

Control mice were handled identically and exposed to the same housing conditions, including placement in the aggressor’s cage with a perforated divider, but without any physical contact or aggression. All behavioral assessments were conducted 30 minutes following the ASDS procedure under standardized conditions.

###### Chronic social defeat stress (CSDS)

Adult C57BL/6J male mice (8−12 weeks old) were subjected to a 10-day chronic social defeat paradigm. Each day, an experimental mouse was introduced into the home cage of an aggressive CD-1 resident mouse for 5 minutes of direct physical confrontation, followed by 24 hours of sensory contact separated by a perforated plexiglass divider. Control mice were housed in similar divided cages with another C57BL/6J mouse, without physical interaction. To ensure consistent stress exposure, experimental mice that exhibited a cumulative attack latency of less than 10 seconds per session were excluded from analysis.

###### 3D behavioral analysis using multi-view motion capture

Three-dimensional behavioral phenotyping was performed using a multi-view kinematic tracking system (BA-DC01, Shenzhen Bayone BioTech). Experimentally male mice (8−12 weeks old) underwent spontaneous exploration assays within a standardized open-field environment, which included: (1) a vertical cylindrical enclosure (height: 30 cm) made of translucent white acrylic polymer, allowing full 360° optical access; (2) an opaque white polycarbonate floor platform (diameter: 40 cm) with surface reflectivity conforming to SIAT behavioral standards; and (3) a modular stainless-steel framework (130 × 130 × 90 cm³) equipped with four Intel RealSense D435 depth cameras positioned orthogonally (90° angular separation; 30 Hz sampling rate), along with an overhead 56-inch LED lighting panel (6500K color temperature) for uniform illumination. Each mouse was introduced into the southeastern quadrant of the arena (Cartesian coordinate: X= +20 cm, Y= −15 cm) under controlled illumination (50 lux) and allowed to acclimate for 15 minutes. Multi-view video capture device and 3D motion-capture system tracked markerless pose estimations of 16 key body parts including the nose, left ear, right ear, neck, left front limb, right front limb, left hind limb, right hind limb, left front claw, right front claw, left hind claw, right hind claw, back, root tail, middle tail, and tip tail.

Prior to each session, spatial calibration was performed using a 25-point reference grid (10-cm spacing), achieving sub-millimeter reconstruction accuracy (<0.5 mm RMS error) via triangulation algorithms. To ensure trial consistency, the arena was sterilized with 75% ethanol and background intensity was normalized between sessions. Raw kinematic data were processed through a series of steps including camera-to-world coordinate transformation, Kalman filter-based noise suppression, and velocity-thresholded event detection (movement onset defined as >2 cm/s). These procedures yielded high-resolution spatiotemporal locomotor trajectories and ethogram classifications, following validated analytical pipelines.^17^

###### Elevated plus maze (EPM) testing

Male mice (8−12 weeks old) were placed at the center of a plus-shaped maze comprising of two open arms (25×5 cm) and two closed arms (25×5×15 cm) under ambient lighting set at 50 lux. Each trial lasted either 3 or 5 minutes, during which video recordings of behavior were obtained and then analyzed using the ANY-maze automated tracking system. Quantified behavioral parameters included time spent in the open arms, the number of entries into open versus closed arms, and total locomotor activity.

###### Open-field test (OFT)

Male mice (8−12 weeks old) were individually placed in a square open-field arena (50×50×50 cm) under uniform illumination (50 lux). Each trial lasted either 3 or 5 minutes, during which video recordings of behavior were obtained and then analyzed using ANY-maze automated tracking software. Behavioral parameters included time spent in the central zone, total distance traveled, and the frequency of rearing behavior. Data were normalized where appropriate to account for variability in overall locomotor activity.

###### Startle reflex testing

Male mice (8−12 weeks old) were secured in a stabilized chamber inside a sound-attenuated startle reflex apparatus (VisuStartle Software XR-XZ208, Shanghai Xinruan, China). Following a 5-minute acclimation period, mice were exposed to randomized acoustic stimuli consisting of 40-ms bursts of 110-dB white noise, interspersed with prepulse tones (20 ms, 65 dB) delivered at variable intervals. Startle amplitude, latency, and prepulse inhibition were recorded and analyzed using VisuStartle software. Data were normalized to baseline startle responses, and statistical significance was assessed using Student’s t-test to evaluate trial effects.

###### Marble-burying test

The test was conducted in a polypropylene mouse cage (42 × 24 × 12 cm) with a metal wire mesh top. The cage floor was layered with 5 cm deep sawdust, on which 20 clean black glass marbles (diameter: 1.5 cm) were evenly placed at uniform intervals. Each male mouse (8−12 weeks old) was placed in this box for 30 minutes. After 30 minutes, the mouse was removed, and the number of marbles buried by the mouse was tallied (counting marbles with at least two-thirds of their surface area covered by bedding material).

###### Tail suspension test (TST)

For the TST, a commercial tail suspension chamber (Bioseb, US) was employed. Male mice (8−12 weeks old) were individually suspended via tail attachment to the force transducer, and their activity was monitored over a 6-minute session. Immobility time was assessed using Anymaze in conjunction with the Bioseb setup.

###### Sucrose preference test (SPT)

Male mice (8−12 weeks old) were single-housed and habituated to two identical drinking bottles (50 ml Falcon tubes with silicone stoppers) for 48 hours; one bottle contained tap water (Bottle A) and the other contained 1% sucrose solution (Bottle B; Sigma, S9378). Bottle positions were counterbalanced across cages to minimize side bias. After 24 hours of water deprivation, baseline fluid consumption was measured during a 4-hour habituation session (09:00–13:00) with both bottles available. Following 24 hours of free access to water, mice underwent a formal 24-hour preference test under identical conditions. Bottle weights (±0.01 g) were measured before and after each phase to calculate fluid consumption, with adjustments made for evaporation using parallel control bottles. Sucrose preference (%) was calculated as: (sucrose intake / (sucrose + water intake)) × 100. Mice exhibiting <60% baseline sucrose preference or technical failures (e.g., bottle leakage >0.5 g) were excluded in accordance with the criteria described by Golden et al.^38^

#### Serum corticosterone and norepinephrine analysis

Blood samples were collected from mice via retro-orbital puncture under isoflurane anesthesia. Serum was obtained by centrifugation at 3000 ×g for 15 minutes and stored at −80°C until further analysis. Circulating levels of corticosterone and norepinephrine were measured using commercial ELISA kits (Abcam, ab108821 and ab287789, respectively), following manufacturer’s instructions. Absorbance was read using a BioTek Synergy H1 microplate reader, and hormone concentrations were normalized to total serum protein content.

#### 9.4T fMRI acquisition from mouse brains *in vivo*

##### Data acquisition

30 min after ASDS, mice were anesthetized with isoflurane (induced at 3% and maintained at 1.5%) and all MRI experiments were performed on a 9.4T scanner with a 30-cm diameter bore (uMR 9.4 T, United Imaging Life Science Instrument, Wuhan, China). An 86-mm diameter transmit volume coil and a Mouse Brain Surface Coil-3 (MBSC3) for RF reception was used. High-resolution structural images were acquired with a Fast Spin Echo T2-weighted (FSE-T2) sequence (TR = 2500 ms, TE = 41.3 ms, FOV = 20 × 20 mm², matrix = 208 × 208, slice thickness = 0.3 mm, no gap). Subsequently, Blood Oxygen Level Dependent (BOLD) functional data were obtained using a gradient-echo echo-planar imaging (GE-EPI) sequence optimized for higher signal-to-noise ratio (TR = 2700 ms, TE = 14.5 ms, FOV = 15 × 19 mm², matrix = 76 × 96, slice thickness = 0.3 mm, no gap).

#### Data analysis

All data processing was performed within MATLAB (The MathWorks, Inc.) using either the SPM12 (Wellcome Centre for Human Neuroimaging, London, UK)^39^ toolbox or custom code. Analysis began by expanding voxels 10-fold, followed by slice timing correction based on slice number, order, and TR. Head motion correction was then applied, with motion parameters (translation < 1 voxel, rotation < 2° across all subjects) estimated for each scan session. Functional MRI (fMRI) data were spatially normalized to a standard template associated with the brain atlas^40^ using the T2-weighted image. Gaussian smoothing was then applied using a smoothing kernel of [4 4 6]. Following this preprocessing pipeline, the DPABI v8.2^41^ toolbox was used to calculate the Amplitude of Low-Frequency Fluctuation (ALFF) parameter maps from the fMRI data. Voxel-wise two-sample t-tests were performed on the ALFF parameter maps using the SPM toolbox. The statistical results were thresholded at a significance level of p < 0.01 with a cluster size > 100 voxels.

#### Assessment of baseline spontaneous neural activity within the targeted CLA

Functional MRI data preprocessing was executed using a combination of AFNI (Analysis of Functional NeuroImages)^42^, ANTs (Advanced Normalization Tools)^43^, and custom MATLAB scripts (MathWorks, Natick, MA). First, slice-timing correction was applied to the functional volumes using AFNI’s 3dTshift to account for the interleaved slice acquisition pattern. Subsequently, head motion correction was performed using AFNI’s 3dvolreg, aligning all time-series volumes to the first volume of the run while simultaneously extracting the six rigid-body motion parameters. For spatial normalization, subject-specific manual transformations were used to ensure anatomical accuracy. These manual transformations, previously defined for each subject using ITK-SNAP^44^, were applied to the motion-corrected functional images using the ANTs toolkit (antsApplyTransforms). The 4D time-series were warped to a study-specific mouse brain template using B-spline interpolation. To mitigate the effects of physiological noise and motion artifacts, a nuisance covariate regression pipeline was applied in MATLAB. The covariates included the demeaned six head motion parameters, the mean signal extracted from an eroded white matter and cerebrospinal fluid (CSF) mask, and the whole-brain global signal. Following regression, spatial smoothing was applied using a 0.3 mm full-width at half-maximum (FWHM) in-plane Gaussian kernel to improve the signal-to-noise ratio.

To evaluate alterations in baseline spontaneous local neural activity, we calculated the mean Amplitude of Low-Frequency Fluctuations (mALFF). The preprocessed time-series data were first linearly detrended to remove scanner signal drift. A Fast Fourier Transform (FFT) was then applied to convert the time-domain signals into the frequency domain, extracting the power spectrum. The Amplitude of Low-Frequency Fluctuations (ALFF) was calculated as the sum of amplitudes within the standard low-frequency band (0.01–0.1 Hz). To account for inter-subject variability in baseline physiological states (e.g., varying depths of anesthesia) and global scanner scaling effects, the raw ALFF values were normalized. Specifically, the ALFF value of each region was divided by the global mean ALFF calculated across all voxels within the whole-brain mask, generating the standardized mALFF metric.

Given the highly specific stereotaxic targeting of the hM3D(Gq) viral injection, our functional analysis utilized a hypothesis-driven, a priori region of interest (ROI) approach to avoid signal dilution from uninfected tissue. The target ROI was anatomically anchored to the spatial coordinates of the injection site in the left voxel neighborhood was generated around the injection center coordinate. To prevent partial volume effects and signal contamination from the adjacent regions, this neighborhood was strictly intersected with the anatomical boundaries of the claustrum defined by the standard mouse atlas. The resulting custom ROI exclusively contained voxels that were both within the anatomical claustrum and directly at the site of chemogenetic manipulation.

Group-level statistical analyses were conducted in MATLAB. Because our experimental design established a strong a priori hypothesis regarding the localized activation of the injected claustrum, statistical testing was restricted exclusively to the custom injection ROI. The mean mALFF values within this target ROI were extracted for both the experimental (3DQ) and control (mCherry) groups. An independent two-sample t-test (two-tailed) was performed to determine the statistical significance of group differences in baseline spontaneous activity. Because the analysis was restricted to a single predefined anatomical target rather than an exploratory whole-brain parcellation, multiple comparison correction was not required. The threshold for statistical significance was set at *p* < 0.05.

#### Single-nucleus RNA sequencing (snRNA-seq)

Tissue Acquisition & Nuclear Isolation: Acquired CLA samples (n = 5 mice/group) were flash-frozen on dry ice. For snRNA-seq, bilateral CLA tissues were homogenized in ice-cold buffer using a Dounce homogenizer (Wheaton, 357542). Homogenates were mixed 1:1 with OptiPrep™ and centrifuged (10,000×g, 20min, 4°C). Pelleted nuclei were washed, resuspended in DMEM/F12+10% FBS, and adjusted to 400 nuclei/μL (hemocytometer). All solutions contained RNase inhibitor (60U/mL). Library Construction & Sequencing: snRNA-seq libraries were generated with Chromium Next GEM Single Cell 3’ Reagent Kits v3.1 (10x Genomics). Briefly, a 40-μL nuclei suspension was combined with RT reagents for droplet encapsulation. Barcoded cDNA was synthesized by PCR and used for library preparation. Library QC involved Qubit™ quantification and Fragment Analyzer™ sizing. Sequencing and Analysis: Using Seurat (v5.2.0), we performed differential gene expression analysis with the FindAllMarkers function, specifying MAST as the statistical test. DEGs were functionally annotated by clusterProfiler (version 4.4.4).

#### Multiplex fluorescence *in situ* hybridization (mulFISH)

We performed mulFISH following protocols adapted from Kunyu Biotech.^45^ *Probe design and hybridization*: Spatially specific targeting probes were custom-designed by Spatial FISH Ltd. Tissue samples fixed in 4% paraformaldehyde were mounted in reaction chambers for subsequent processing. After methanol-mediated dehydration and denaturation, hybridization buffer containing the targeting probes was applied, and samples were incubated overnight at 37°C. *Ligation and amplification*: Following three washes in PBST, ligation was carried out using a ligation mix at 25°C for 3 hours. The samples were then washed and subjected to rolling circle amplification with Phi29 DNA polymerase at 30°C overnight. *Detection and imaging*: Fluorescent detection probes complementary to the amplified sequences were hybridized in hybridization buffer. After final ethanol dehydration, samples were mounted for imaging. Spatial localization of RNA transcripts was decoded based on fluorescent puncta captured using a Leica THUNDER Imaging System (20× objective, NA = 0.80).

#### Chemogenetic manipulation and stereotactic virus injection

Adeno-associated viruses (AAVs), AAV-DIO-hM3Dq-mCherry, AAV-DIO-hM4Di-mCherry and AAV-DIO-mCherry were packaged by Brain Case (Shenzhen, China) at titers of 5×10^12^ particles per ml. Adult male Gad2-Cre or Egr2-Cre mice (8−12 weeks old) were deeply anesthetized with sodium pentobarbital (1%, 10 mL/kg body weight, i.p.; Sigma-Aldrich, #P3761) and positioned in a stereotaxic instrument (RWD Life Science Inc., Shenzhen, China) with their heads fixed. Virus was delivered using a microinjector pump (UMP3/Micro4, World Precision Instruments, USA) connected to a 10-μL Hamilton syringe, at an injection rate of 50 nL/min. After infusion, the injection needle was left in place for 10 minutes to minimize reflux. For DREADD-based manipulations, 150 nL of virus was bilaterally injected into the Claustrum (CLA) using the following coordinates: anterior-posterior (AP) +1.00 mm, mediolateral (ML) ± 2.80 mm, dorsoventral (DV) − 3.70 mm. To activate hM3Dq-expressing neurons, clozapine-N-oxide (CNO; Med Chem Express, #HY-17366) was administered intraperitoneally (1 mg/kg). Behavioral testing was performed within a 60-minute window, starting 30 minutes post-injection. All behavioral experiments were conducted under ambient lighting (∼200 lux).

In male Gad2-Cre (8−12 weeks old) mice used for circuit-specific manipulation and retrograde labeling of ACC-projecting CLA^Gad2⁺^ neurons, a viral cocktail of AAV-DIO-EGFP-T2A-TVA and AAV-DIO-N2cG was injected into the CLA using the coordinates given above. Concurrently, AAV-fDIO-hM3Dq-GFP (or control AAV-fDIO-GFP) was delivered into the ACC (AP +1 mm, ML ±0.3 mm, DV −1.8 mm). For targeted viral delivery to claustrum Gad2⁺ neurons, the viral titer was adjusted to below 10¹² v.g./mL, and the injection volume was set to 80 nL. Viruses were injected at a slow rate of 20 nL/min, followed by a post-injection dwell time of more than 10 min to minimize viral spread to adjacent brain regions and ensure specific targeting of the claustrum.

Three days prior to behavioral testing, EnvA-pseudotyped, glycoprotein-deleted rabies virus (RV-EnvA-ΔG-mCherry-Flp) was injected into the CLA for retrograde labeling of ACC→CLA^Gad2⁺^ projections. Chemogenetic activation of this circuit was achieved by daily CNO administration (1 mg/kg, i.p.) over seven consecutive days, during which behavioral outcomes were quantitatively assessed.

#### Optogenetic stimulation

Stereotaxic surgery was performed on adult Egr2-Cre mice (8−12 weeks, male) with bilateral injections of AAV-DIO-ChR2-GFP (150 nL/site; 5×10^12^ vg/ml, Brain Case) into the CLA. Concurrently, 200-μm optical fibers (0.37 NA) were implanted above the ACC (coordinates: AP +1.0 mm, ML ±0.3 mm, DV −1.5 mm) to enable terminal stimulation of CLA-originating Egr2 projections. Mice were given 4 weeks to recover and to ensure sufficient transgene expression.

To minimize stress and facilitate behavioral testing, animals were habituated to fiber attachment in their home cages for 3 consecutive days (15 min/day in home cages). During behavioral testing, blue light stimulation (470 nm, 10 Hz, 40-ms pulse width, 5 mW) was delivered through the optic cannula during the designated “light-on” periods. A control group injected with AAV-DIO-GFP underwent the same surgical procedures and received identical light stimulation.

#### Immunohistochemistry

Mice were euthanized via overdose of 1% sodium pentobarbital (15 mL/kg body weight, i.p.) and transcardially perfused with phosphate-buffered saline (PBS), followed by 4% paraformaldehyde (PFA; Aladdin, #C104188) in PBS. Brains were extracted, post-fixed overnight in 4% PFA at 4°C, and cryoprotected in 20% sucrose for 24 hours followed by 30% sucrose for 48 hours. Tissue was embedded in O.C.T. compound (Tissue-Tek® Optimal Cutting Temperature) and sectioned coronally at 40-μm thickness using a cryostat microtome (Leica CM1950, Germany).

Sections were washed three times in PBS (3 min per wash, room temperature) to remove residual O.C.T., then blocked in PBS containing 0.3% Triton X-100 and 3% bovine serum albumin (BSA) for 1 hour at room temperature. Primary antibodies were incubated overnight at 4°C in dilutions of 1:200-1:300 in PBS containing 0.1% Triton X-100 and 3% BSA. Sections were subsequently incubated for 2 hours at room temperature with secondary antibodies: Alexa Fluor® 405-, 488-, 594-, or 647-conjugated goat anti-rabbit or anti-rat IgG (1:200-1:300, Invitrogen). After mounting with DAPI-containing anti-fade reagent (ProLong™ Gold, Life Technologies), sections were imaged using an Olympus VS120-S6-W slide scanner or Zeiss LSM 980 confocal microscope. Neuroanatomical regions were identified according to The Mouse Brain in Stereotaxic Coordinates (Franklin and Paxinos, 1997).

#### Patch-clamp electrophysiology

##### Combined patch-clamp and optogenetics

We employed combined patch-clamp/optogenetics to record action potential dynamics in excitatory neurons during blue light (470 nm) activation of ChR2-expressing Gad2 neurons. Three weeks after stereotaxic injection of AAV-DIO-ChR2-GFP into the CLA of Gad2-Cre mice, coronal CLA slices (250 μm; bregma +1.4 to +0.6 mm) were prepared in ice-cold modified ACSF (mM: 205 Sucrose 2.5 KCl, 1.25 NaHPO₄, 25 NaHCO, 10 Glucose, 0.5 CaCl, 7.5 MgCl₂) using a Leica VT-1200S vibrotome. Slices were recovered after 30 min at 32−34°C in oxygenated standard ACSF (mM: 125 NaCl, 2.5 KCl, 1.3 NaH₂PO₄, 25 NaHCO₃, 25 Glucose, 2 CaCl₂, 1 MgCl₂; pH 7.3−7.4), then held at room temperature. All solutions: were between 300−320 mOsm/kg. All drugs used were from Sigma-Aldrich. Slices were visualized under Nikon FN1 microscope with IR optics. Recording used oxygenated ACSF perfusion (2 mL/min, RT). Pipettes (5−10 MΩ; Sutter P-97) contained (mM): 125 K-gluconate, 20 KCl, 0.5 EGTA, 10 HEPES, 10 Na2-creatine, 4 Mg-ATP, 0.3 Na-GTP. AP firing was evoked by 2-s depolarizing steps (current-clamp). Signals amplified via HEKA EPC-10, filtered at 2.9 kHz and 10 kHz, digitized at 20 kHz (Patchmaster). Series resistance (10-30 MΩ) was monitored; data discarded if >30% change occurred. Optical activation of Gad2^+^ neurons via blue light irradiation confirmed ChR2 functionality whilst electrophysiological responses of surrounding excitatory neurons were recorded.

##### Patch-Clamp in Gad2^+^ neuron in CLA

Method guided by Jarvis Biological Pharmaceutical Co., Ltd (Wuhan, China). 7 days after ASDS or CSDS, mice were anesthetized with sodium pentobarbital (45 mg/kg) via intraperitoneal injection (i.p.) and intracardially perfused with ice-cold slicing solution (in mM, 209 sucrose, 3.1 sodium pyruvate, 22 glucose, 1.25 NaH₂PO₄, 12 sodium l-ascorbate, 4.9 MgSO₄·7H₂O, 26 NaHCO₃; 95% O₂/5% CO₂, pH 7.2–7.4). Coronal slices (300 μm) of the claustrum (CLA) were prepared using a vibratome (Leica VT1200 S, Leica Biosystems) and incubated in aCSF (in mM, 128 NaCl, 3 KCl, 24 NaHCO₃, 2 MgCl₂, 1.25 NaH₂PO₄, 10 d-glucose, 2 CaCl₂; oxygenated, pH 7.2–7.4, 295–305 mOsm) at 28°C for 1 hour, then equilibrated at room temperature for patch-clamp recordings. Neurons were voltage-clamped at −70 mV in voltage-clamp mode to record spontaneous excitatory postsynaptic currents (sEPSC). sEPSC pipettes (4–6 MΩ) were filled with an internal solution comprising (in mM, 122.5 Cs-gluconate, 17.5 CsCl, 0.2 EGTA, 10 HEPES, 1 MgCl₂, 4 Mg-ATP, 0.3 Na-GTP, 5 QX314, pH 7.25, 280–300 mOsm), with 100 μM PTX in the bath. For action potentials, current-clamp mode used pipette solution (in mM, 140 K-gluconate, 3 KCl, 2 MgCl₂, 10 HEPES, 0.2 EGTA, 2 Na₂ATP, 285–295 mOsm, pH 7.2). All data was obtained by pCLAMP 10 (Axon Instruments, Molecular Devices, San Jose, CA) and the MultiClamp 700B amplifier (Molecular Devices, Sunnyvale, CA). For the AP and sEPSC recordings analyzed, signals were low-pass filtered at 2 kHz and digitized at a sampling rate of 20 kHz. (Molecular Devices, Jarvisbio, Wuhan, China).

##### Patch-Clamp in Egr2^+^ neuron in CLA

Male Egr2-Cre mice (provided by the client) were injected with AAV-DIO-mCherry, housed in Jiawei Animal Experiment Platform for 4 weeks, and divided into Control, ASDS, and CSDS groups. Electrophysiological recordings used MultiClamp 700B amplifier, VT1200S vibratome, BX51WI upright microscope, Sutter P-1000 puller, and 1550A digitizer; data were collected via Clampex 10 and analyzed with Clampfit and GraphPad Prism. ACSF (95% O₂ + 5% CO₂) with 100 μM PTX (for sEPSC) was circulated. Brain slices were transferred to the recording chamber; 4–6 MΩ electrodes filled with internal solution were used to form >1 GΩ seals on mCherry-positive CLA neurons. sEPSCs (voltage-clamp, −70 mV) and APs (current-clamp) were recorded for 5 min per neuron.

#### Fiber photometry calcium signals recording

Three weeks following bilateral injections of AAV-DIO-ChrimsonR-mCherry (150 nl/site; 5×10¹² vg/ml, BrainVTA) and AAV-vGluT1-GCaMP6s (150 nL/site; 5×10¹² vg/mL, BrainVTA) into the CLA, Gad2-Cre mice (8–12 weeks, male) were habituated to fiber patch cords attachment in their home cages (≥15 min/day for 3 days) prior to testing. Calcium signals were recorded using a dual-wavelength fiber photometry system (ThinkerTech, Nanjing) equipped with 470-nm and 410-nm^46^ LEDs for GCaMP6s excitation and motion artifact correction, respectively. Both excitation channels were coupled via a single objective to a 200-μm, 0.37 numerical aperture optical fiber (RWD Life Science). Simultaneously, a 590-nm laser (ThinkerTech) was used to activate ChrimsonR-expressing CLA^Gad2⁺^ neurons while recording GCaMP6s signals from CLA^vGluT1⁺^ neurons. Light intensity at the fiber tip was maintained at 25–30 μW. GCaMP6s fluorescence emission was collected through the same optical path, separated by a dichroic mirror, and detected by a photodetector. Calcium signal processing and analysis were performed using custom scripts in MATLAB R2017b.

For ACC calcium signal recordings, test mice were placed in a cylindrical wire enclosure, and aggressive CD1 mice were allowed to freely explore and interact in the surrounding open field. Time zero was aligned to the moment when the CD1 mouse approached within 2 cm of the enclosure.

#### Statistical summary

Unless otherwise specified, comparisons between two groups were performed using unpaired two-tailed Student’s t-tests. Comparisons involving four groups were analyzed using two-way analysis of variance (ANOVA). Where significant main effects or interactions were found by two-way ANOVA, post hoc pairwise comparisons between groups were performed using Tukey’s multiple comparisons test. All statistical analyses were conducted using GraphPad Prism version 9.0 (GraphPad Software, San Diego, CA). Data are presented as mean ± standard error of the mean (SEM). Statistical significance was defined as *P* < 0.05.

## Supporting information

Manuscript

## Acknowledgments

We appreciate the very helpful suggestions and comments on earlier versions of the manuscript by Prof. Minmin Luo and Prof. Xiang Yu. We are grateful to Prof. Teng Chen and Prof. Xinshe Liu for their valuable support. The 3D behavioral system and related data analysis were supported by Prof. Wang Feng. We thank Yingying Du and Tong Ye for their contributions to the MRI data acquired and analysis. We thank Zili Liu for her technical support for the patch-clamp data acquired. The technical support for electrophysiology from Jarvis Biological Pharmaceutical Co., Ltd (Wuhan, China) is sincerely appreciated. The figures in this paper were created with BioRender.com.

This research was also supported by the following platforms: Guangdong Provincial Key Laboratory of Brain Connectome and Behavior (2023B1212060055); Shenzhen Brain Science Infrastructure; SIAT-HKUST Joint Laboratory for Brain Science; Shenzhen Science and Technology Program, Shenzhen Key Laboratory of Neuroimmunomodulation for Neurological Diseases (ZDSYS20220304163558001).

## Funding

This study was supported by the Major Program of National Natural Science Foundation of China (T2394532 F.Y.), the Shenzhen Medical Research Fund (B2402018 J.T., B2302011 F.Y., E250200313 F.Y.), the National Natural Science Foundation of China (32371070 J.T.), the National Key R&D Program of China (No.2024YFC3406700 J.T.), CAS Project for Young Scientists in Basic Research (Grant No. YSBR-126), the Key-Area Research and Development Program of Guangdong Province (2023B0303040004 J.T.), and the Shenzhen Science and Technology Program (JCYJ20220818101615033 D.L., JCYJ20210324101813035 D.L., JCYJ20241202125015020 Q.X., 202208183000430 F.Y.). The Natural Science Foundation of Shanghai (24ZR1451500, Z.M.). Shanghai Scientific Instruments and Chemical Reagents Project (24142201100, Z.M.).

## Author contributions

J.T. conceived the study. D.L., Z-J.L., G-J. S., X-X. Z, H-H H and J.S. performed the experiments. J.T., D.L., G-J. S., Z-J.L., Q.X., Y-Y. Q, Z-W. M and S.C. analyzed the data. L-P. W., F.Y. and Y.C. provided suggestions on the manuscript. D.L. and J.T. wrote the manuscript. J.T. supervised the project.

## Declaration of interests

The authors have declared that no conflict of interest exists.

## Notes

### Competing Interest Statement

The authors have declared no competing interest.

### Summary of Updates

Based on the reviewers' comments, we have modified the specific promoters and the acute and chronic stress models, as well as other relevant aspects.

