## Supplementary material for "Claustrum–ACC reciprocal circuits modulate stress responses across acute and chronic states": Manuscript

Figure S1

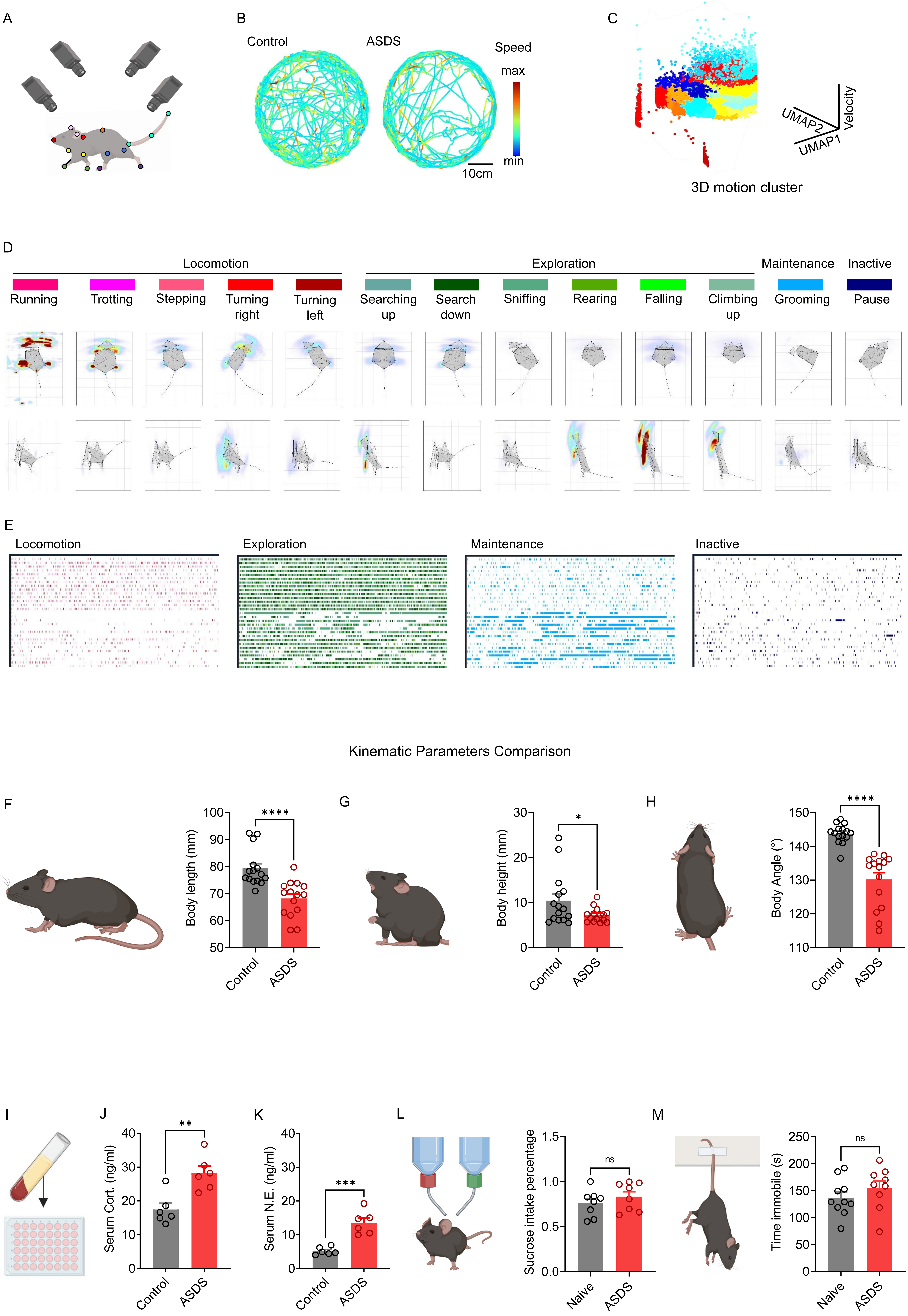

A
B
Control
ASDS
Speed
max
min
10cm
C
UMAP2
UMAP1
Velocity
3D motion cluster
D
Locomotion
Exploration
Maintenance
Inactive
Running
Trotting
Stepping
Turning right
Turning left
Searching up
Search down
Sniffing
Rearing
Falling
Climbing up
Grooming
Pause
E
Locomotion
Exploration
Maintenance
Inactive
Kinematic Parameters Comparison
F
Body length (mm)
****
Control
ASDS
G
Body height (mm)
*
Control
ASDS
H
Body Angle (°)
****
Control
ASDS
I
J
Serum Cort. (ng/ml)
**
Control
ASDS
K
Serum N.E. (ng/ml)
***
Control
ASDS
L
Sucrose intake percentage
ns
Naive
ASDS
M
Time immobile (s)
ns
Naive
ASDS

**Figure S1.**

**Acute social defeat stress (ASDS) induces anxiety-like behaviors without depressive phenotypes**

(A) Schematic showing the multi-camera 3D pose reconstruction system used for fine-grained behavioral quantification, with tracked anatomical landmarks (nose, ears, neck, back, tail segments, and limb/claw positions).

(B) Representative 3D movement trajectories of control (left) and ASDS (right) mice, color-coded by instantaneous speed (blue, low; red, high), illustrating altered locomotor patterns after ASDS.

(C) UMAP embedding of 3D behavioral features, colored by velocity, showing separation of movement dynamics between control and ASDS groups.

(D) Behavioral ethogram and representative 3D pose heatmaps, with behaviors grouped into four domains: locomotion (running, trotting, stepping, turning left/right), exploration (searching up/down, sniffing, rearing, falling, climbing up), maintenance (grooming), and inactive (pause).

(E) Temporal distribution of behavioral states in control (top) and ASDS (bottom) mice, revealing reduced exploratory behavior and increased maintenance/inactive states in the ASDS group.

(F–H) Kinematic parameters in ASDS and control mice (n =15/group). (F) Body length ($t_{(28)} = 4.565$, $P < 0.0001$).

(G) Body height ($t_{(28)}$ = 2.104, $P$ = 0.0445). (H) Body angle ($t_{(28)}$ = 6.275, $P$ < 0.0001). Statistical analysis was performed using an unpaired two-tailed Student's *t*-test.

(I–K) Serum stress hormone levels measured by enzyme-linked immunosorbent assay (ELISA) in ASDS and control groups (n=6/group): (I) schematic illustration showing the ELISA workflow, (J) corticosterone (Cort.) levels ($t_{(10)}$ = 3.863, $P$ = 0.0031) and (K) norepinephrine (N.E.) levels ($t_{(10)}$ = 5.598, $P$ = 0.0002). Statistical analysis was performed using an unpaired two-tailed Student's *t*-test.

(L–M) Assessment of depressive-like behaviors (n = 8/group): (L) sucrose preference ratio was unchanged between control and ASDS groups ($t_{(14)}$ = 0.9821, $P$ = 0.3427) and (M) immobility time ($t_{(14)}$ = 1.064, $P$ = 0.3023) in the tail suspension test, showing no significant differences between groups. Statistical analysis was performed using an unpaired two-tailed Student's *t*-test.

Data are presented as mean ± SEM. *$P$ < 0.05, **$P$ < 0.01, ***$P$ < 0.001, ****$P$ < 0.0001, n.s. represents $P$ > 0.05.

Figure S2

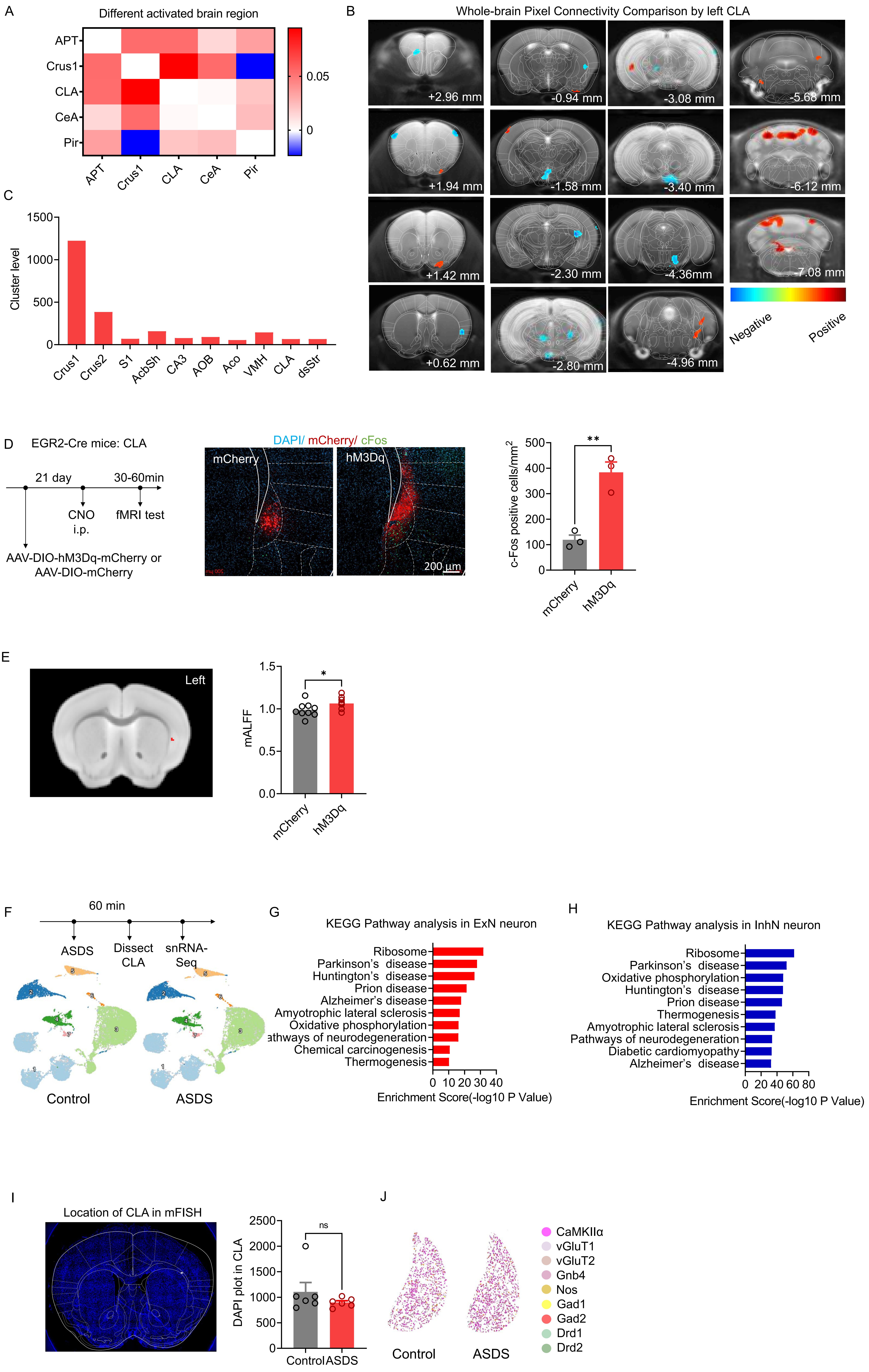

A
Different activated brain region
APT
Crus1
CLA
CeA
Pir
0.05
0
B
Whole-brain Pixel Connectivity Comparison by left CLA
+2.96 mm
-0.94 mm
-3.08 mm
-5.68 mm
+1.94 mm
-1.58 mm
-3.40 mm
-6.12 mm
+1.42 mm
-2.30 mm
-4.36mm
-7.08 mm
+0.62 mm
-2.80 mm
-4.96 mm
Negative
Positive
C
Cluster level
1500
1000
500
0
Crus1
Crus2
S1
AcbSh
CA3
AOB
Aco
VMH
CLA
dsStr
D
EGR2-Cre mice: CLA
21 day
30-60min
CNO
i.p.
fMRI test
AAV-DIO-hM3Dq-mCherry or
AAV-DIO-mCherry
DAPI/ mCherry/ cFos
mCherry
hM3Dq
200 μm
c-Fos positive cells/mm2
500
400
300
200
100
0
**
mCherry
hM3Dq
E
Left
mALFF
1.5
1.0
0.5
0.0
*
mCherry
hM3Dq
F
60 min
ASDS
Dissect
CLA
snRNA-
Seq
Control
ASDS
G
KEGG Pathway analysis in ExN neuron
Ribosome
Parkinson's disease
Huntington's disease
Prion disease
Alzheimer's disease
Amyotrophic lateral sclerosis
Oxidative phosphorylation
athways of neurodegeneration
Chemical carcinogenesis
Thermogenesis
0 10 20 30 40
Enrichment Score(-log10 P Value)
H
KEGG Pathway analysis in InhN neuron
Ribosome
Parkinson's disease
Oxidative phosphorylation
Huntington's disease
Prion disease
Thermogenesis
Amyotrophic lateral sclerosis
Pathways of neurodegeneration
Diabetic cardiomyopathy
Alzheimer's disease
0 20 40 60 80
Enrichment Score(-log10 P Value)
I
Location of CLA in mFISH
DAPI plot in CLA
2500
2000
1500
1000
500
0
ns
Control
ASDS
J
Control
ASDS
CaMKIIα
vGluT1
vGluT2
Gnb4
Nos
Gad1
Gad2
Drd1
Drd2

**Figure S2**

**ASDS engages the CLA as a central hub with altered connectivity and transcriptional responses**

(A–C) CLA-centered functional connectivity changes following ASDS. (A) Correlation analysis among differentially activated brain regions identified by fMRI. (B) Brain-wide map of regions functionally correlated with left CLA activation. (C) Quantification of pixel counts in regions showing significant correlation with the left CLA.

(D–E) Positive control of CLA activation detected by MRI. (D) Targeted activation of EGR2-positive neurons in the CLA, c-Fos density was analyzed using an unpaired two-tailed Student's *t*-test ($t_{(4)} = 5.926$, $P = 0.0041$). (E) Assessment of baseline spontaneous neural activity within the targeted claustrum (CLA). Left: Representative coronal MRI template overlaid with the hypothesis-driven, custom-defined region of interest (ROI) mask (red), localizing the specific site of viral injection within the claustrum. Right: Quantification of the mean Amplitude of Low-Frequency Fluctuations (mALFF) within the injected CLA ROI. Chemogenetic stimulation in the hM3Dq group induced a significant increase in local mALFF compared to the mCherry control group. Circles represent individual subjects (mCherry, n = 9; hM3Dq, n = 10). Bars represent group means. Statistical significance was determined using an independent two-sample t-test ($t_{(17)} = 2.139$, $P = 0.0473$); $*P < 0.05$.

(F–H) Single-nucleus RNA sequencing and KEGG analysis in ASDS and control mice.

(F) Experimental workflow and UMAP visualization of cell-type clustering in control (left) and ASDS (right) mice (ASDS →60 min rest → CLA dissection→ snRNA-seq, n = 2/group from 10 mice), where colors denote distinct cell-type clusters. (G–H) KEGG pathway enrichment analysis of excitatory and inhibitory neurons.

(I–J) Cell-type marker expression in the CLA assessed by multiplex *in situ* hybridization. (I) Region of interest and quantification of DAPI-positive cells, showing no change in cell number. (J) Schematic diagram showing the detected gene markers across ASDS and control groups.

Data are presented as mean ± SEM. $^{*}P < 0.05$, $^{**}P < 0.01$, $^{***}P < 0.001$, $^{****}P < 0.0001$, n.s. represents $P > 0.05$.

Figure S3

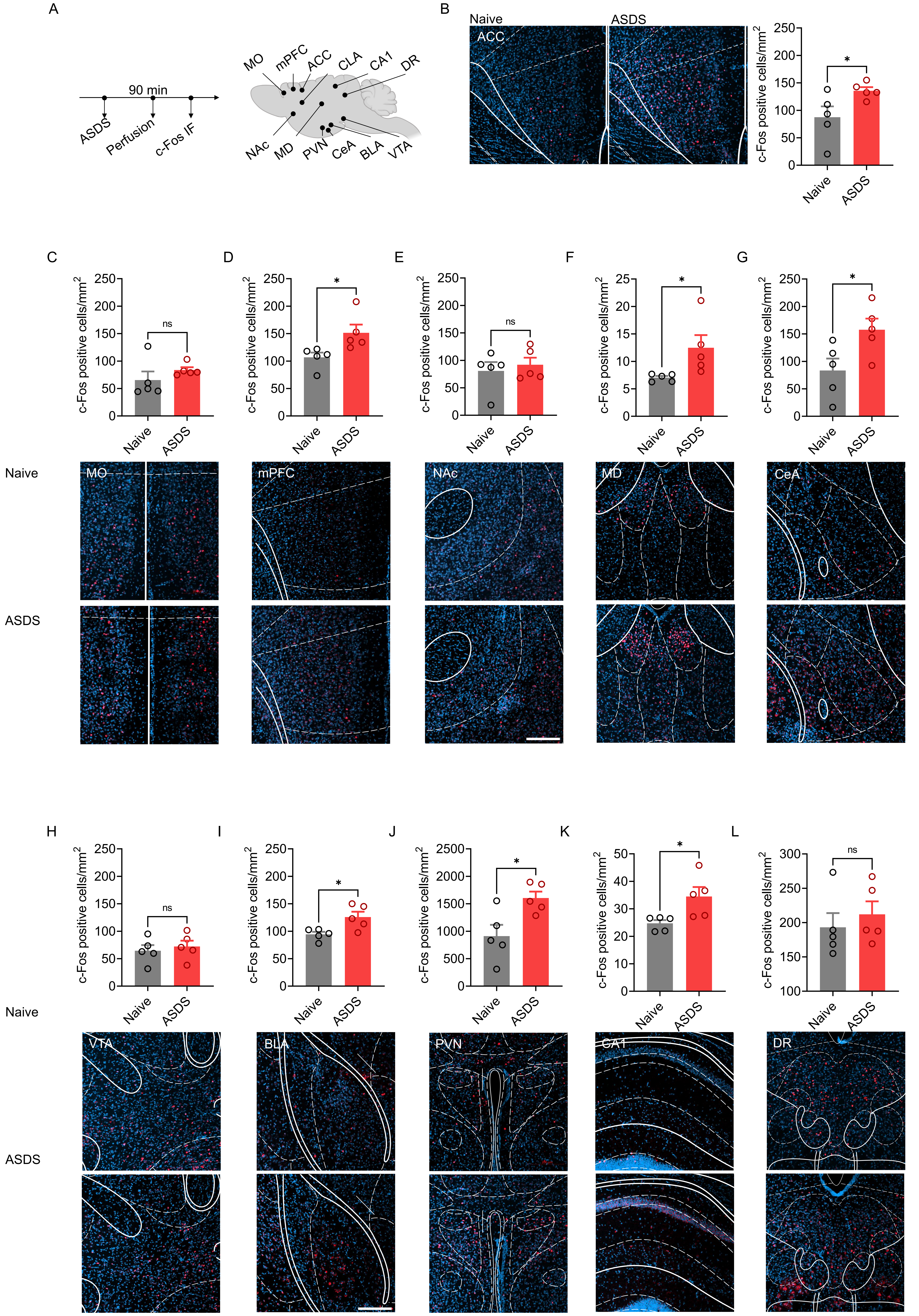

A
90 min
ASDS
Perfusion
c-Fos IF
MO
mPFC
ACC
CLA
CA1
DR
NAc
MD
PVN
CeA
BLA
VTA
B
Naive
ASDS
ACC
c-Fos positive cells/mm²
C
D
E
F
G
ns
*
Naive
ASDS
MO
mPFC
NAc
MD
CeA
H
I
J
K
L
VTA
BLA
PVN
CA1
DR

**Figure S3**

**c-Fos mapping reveals region-specific activation of stress-related brain areas following ASDS**

(A) Schematic diagram showing c-Fos screening in stress-related brain regions.

(B–L) Quantification of c-Fos expression across brain regions in control and ASDS mice (n = 5/group). (B) Schematic diagram showing c-Fos in the ACC region, analyzed using an unpaired two-tailed Student's *t*-test ($t_{(8)}$ = 2.323, $P$ = 0.0487). (C) Schematic diagram showing c-Fos in the ACC region, analyzed using an unpaired two-tailed Student's *t*-test ($t_{(8)}$ =1.115, $P$ = 0.2972).n(D) Schematic diagram showing c-Fos in the mPFC region, analyzed using an unpaired two-tailed Student's *t*-test ($t_{(8)}$ = 2.603, $P$ = 0.0315). (E) Schematic diagram showing c-Fos in the NAc region, analyzed using an unpaired two-tailed Student's *t*-test ($t_{(8)}$ = 0.5500, $P$ = 0.5973). (F) Schematic diagram showing c-Fos in the MD region, analyzed using an unpaired two-tailed Student's *t*-test ($t_{(8)}$ = 2.352, $P$ = 0.0465).

(G) Schematic diagram showing c-Fos in the CeA region, analyzed using an unpaired two-tailed Student's *t*-test ($t_{(8)}$ = 2.485, $P$ = 0.0378). (H) Schematic diagram showing c-Fos in the VTA region, analyzed using an unpaired two-tailed Student's *t*-test ($t_{(8)}$ = 0.5256, $P$ = 0.6134). (I) Schematic diagram showing c-Fos in the BLA region, analyzed using an unpaired two-tailed Student's *t*-test ($t_{(8)}$ = 2.906, $P$ = 0.0197). (J) Schematic diagram showing c-Fos in the PVN region, analyzed using an unpaired two-tailed Student's *t*-test ($t_{(8)}$ = 2.934, $P$ = 0.0189).

(K) Schematic diagram showing c-Fos in the CA1 region, analyzed using an unpaired two-tailed Student's *t*-test ($t_{(8)}$ = 2.701, $P$ = 0.0270).

(L) Schematic diagram showing c-Fos in the DRN region, analyzed using an unpaired two-tailed Student's *t*-test ($t_{(8)}$ = 0.6712, $P$ = 0.5210).

Data are presented as mean ± SEM. *$P$ < 0.05, **$P$ < 0.01, ***$P$ < 0.001, ****$P$ < 0.0001, n.s. represents $P$ > 0.05.

Figure S4

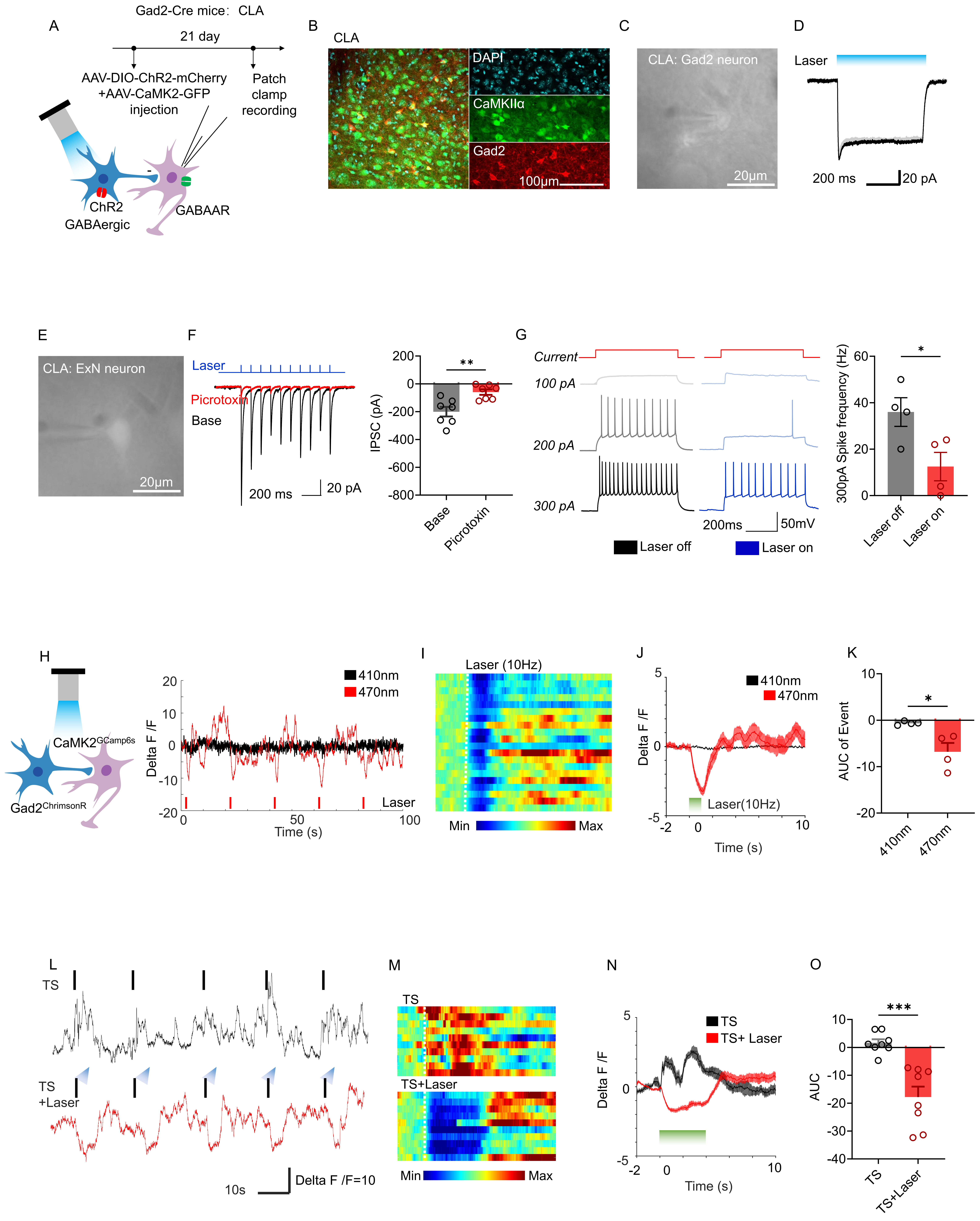
A
Gad2-Cre mice: CLA
21 day
AAV-DIO-ChR2-mCherry
+AAV-CaMK2-GFP
injection
Patch
clamp
recording
ChR2
GABAergic
GABAAR
B
CLA
DAPI
CaMKIIα
Gad2
100μm
C
CLA: Gad2 neuron
20μm
D
Laser
200 ms
20 pA
E
CLA: ExN neuron
20μm
F
Laser
Picrotoxin
Base
200 ms
20 pA
IPSC (pA)
**
Base
Picrotoxin
G
Current
100 pA
200 pA
300 pA
200ms
50mV
Laser off
Laser on
300pA Spike frequency (Hz)
*
Laser off
Laser on
H
CaMK2GCamp6s
Gad2ChrimsonR
410nm
470nm
Delta F /F
Time (s)
Laser
I
Laser (10Hz)
Min
Max
J
410nm
470nm
Delta F /F
Laser(10Hz)
Time (s)
K
AUC of Event
*
410nm
470nm
L
TS
TS
+Laser
10s
Delta F /F=10
M
TS
TS+Laser
Min
Max
N
TS
TS+ Laser
Delta F /F
Time (s)
O
***
AUC
TS
TS+Laser

**Figure S4**

**GABAergic CLA$^{Gad2}$ neurons inhibit claustral excitatory neurons and constrain stress-induced neural activity**

(A–D) Optogenetic and electrophysiological validation of CLA$^{Gad2}$ inhibitory neurons. (A) Experimental workflow for optogenetic manipulation and electrophysiology. (B) Immunostaining of Gad2+ and CaMKIIα+ neurons in CLA. (C) Targeted patch-clamp recording from a Gad2 neuron. (D) Representative light-evoked inhibitory postsynaptic current (IPSC).

(E–G) CLA$^{Gad2}$ neurons suppress excitatory neuronal activity in the CLA. (E) Targeted recording from claustral excitatory neurons (ExN). (F) Light-evoked IPSCs are blocked by picrotoxin, analyzed using an unpaired two-tailed Student's *t*-test (n = 7/group, $t_{(12)} = 3.556$, $P = 0.0040$). (G) Optogenetic activation of CLA$^{Gad2}$ neurons suppresses excitatory neuronal spiking, analyzed using an unpaired two-tailed Student's *t*-test (n = 4/group, $t_{(6)} = 2.703$, $P = 0.0354$).

(H–K) Optical activation of CLA$^{Gad2}$ neurons suppresses overall claustral neural activity in vivo. (H) Calcium signal time course under control and laser conditions. (I) Heatmaps of single-neuron activity. (J) Averaged $\Delta F/F$ traces. (K) Quantification of AUC showing reduced activity following activation, analyzed using an unpaired two-tailed Student's *t*-test (n = 4/group, $t_{(6)} = 3.233$, $P = 0.0178$).

(L–O) CLA$^{Gad2}$ activation suppresses stress-induced excitatory activity. (L) Calcium signal time course during tail suspension (TS) with and without laser stimulation. (M)

Heatmaps showing reduced neural activity under TS + laser. (N) Averaged *ΔF/F* traces. (O) AUC quantification showing suppression of stress-evoked activity, analyzed using an unpaired two-tailed Student's *t*-test (n = 8/group, $t_{(14)} = 4.902$, $P = 0.0002$). Data are presented as mean ± SEM. *$P < 0.05$, **$P < 0.01$, ***$P < 0.001$, ****$P < 0.0001$, n.s. represents $P > 0.05$.

Figure S5

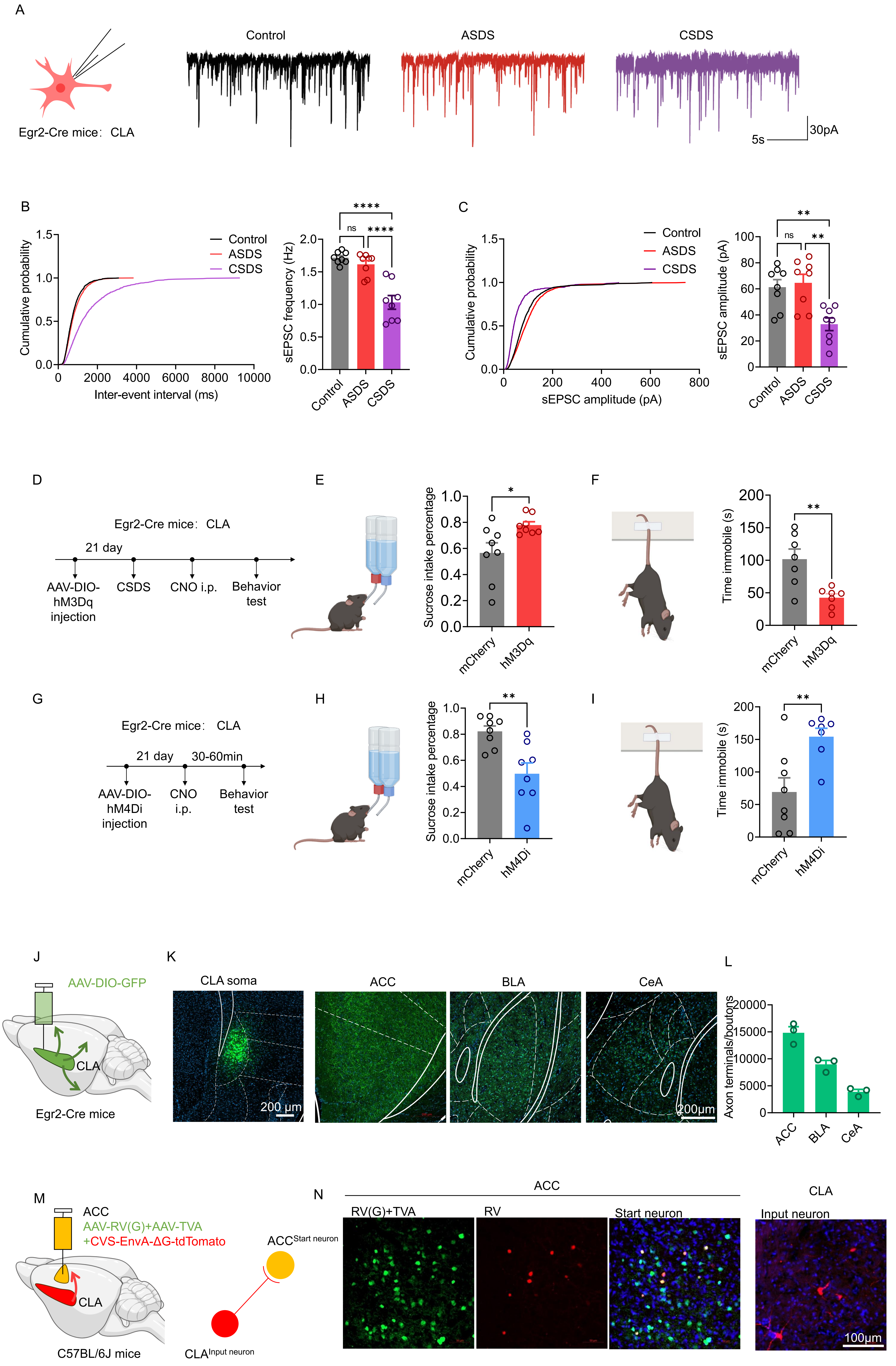
A
Egr2-Cre mice：CLA
Control
ASDS
CSDS
5s
30pA
B
Cumulative probability
Inter-event interval (ms)
Control
ASDS
CSDS
sEPSC frequency (Hz)
ns
****
C
Cumulative probability
sEPSC amplitude (pA)
sEPSC amplitude (pA)
**
D
Egr2-Cre mice：CLA
21 day
AAV-DIO-hM3Dq injection
CSDS
CNO i.p.
Behavior test
E
Sucrose intake percentage
mCherry
hM3Dq
*
F
Time immobile (s)
**
G
Egr2-Cre mice：CLA
21 day
30-60min
AAV-DIO-hM4Di injection
CNO i.p.
Behavior test
H
Sucrose intake percentage
mCherry
hM4Di
I
Time immobile (s)
J
AAV-DIO-GFP
CLA
Egr2-Cre mice
K
CLA soma
ACC
BLA
CeA
200 μm
200μm
L
Axon terminals/boutons
ACC
BLA
CeA
M
ACC
AAV-RV(G)+AAV-TVA
+CVS-EnvA-ΔG-tdTomato
CLA
C57BL/6J mice
ACC Start neuron
CLA Input neuron
N
ACC
RV(G)+TVA
RV
Start neuron
CLA
Input neuron
100μm

**Figure S5**

**CLA$^{Egr2}$ neurons bidirectionally regulate depressive-like behaviors and define claustral projection architecture**

(A–C) Chronic stress reduces excitatory synaptic input onto CLA$^{Egr2}$ neurons (n = 8/group). (A) Representative traces of sEPSCs recorded from Egr2-positive CLA neurons in control, ASDS, and CSDS groups. (B) Cumulative probability distributions and quantification of sEPSC inter-event intervals and frequency (one-way ANOVA, $F_{(2, 21)} = 25.84$, $P < 0.0001$). (C) Cumulative distributions and quantification of sEPSC amplitude (one-way ANOVA, $F_{(2, 21)} = 9.117$, $P = 0.0014$).

(D–F) Chemogenetic activation of CLA$^{Egr2}$ neurons alleviates depressive-like behaviors following CSDS (n = 8/group). (D) Experimental workflow for chemogenetic manipulation of CLA$^{Egr2}$ neurons in Egr2-Cre mice. (E) Increased sucrose preference, analyzed using an unpaired two-tailed Student's *t*-test ($t_{(14)} = 2.601$, $P = 0.0209$). (F) Reduced immobility time in the tail suspension test, analyzed using an unpaired two-tailed Student's *t*-test ($t_{(14)} = 3.573$, $P = 0.0038$).

(G–I) Chemogenetic inhibition of CLA$^{Egr2}$ neurons induces depressive-like behaviors (n = 8/group). (G) Experimental workflow for chemogenetic inhibition of CLA$^{Egr2}$ neurons. (H) Decreased sucrose preference, analyzed using an unpaired two-tailed Student's *t*-test ($t_{(14)} = 3.487$, $P = 0.0036$). (I) Increased immobility time in the tail suspension test, analyzed using an unpaired two-tailed Student's *t*-test ($t_{(13)} = 3.223$, $P = 0.0067$).

(J–N) Anatomical organization of claustral excitatory projections and inputs. (J) Anterograde tracing of CLA excitatory neuron projections. (K) Representative images showing axonal terminals in ACC, CeA, BLA, and CeL. (L) Quantification of projection density, indicating strong innervation of the ACC and CeA. (M) Monosynaptic retrograde tracing from the ACC. (N) Representative images confirming monosynaptic input from ACC to CLA.

Data are presented as mean ± SEM. $*P < 0.05$, $**P < 0.01$, $***P < 0.001$, $****P < 0.0001$, n.s. represents $P > 0.05$.
